# Defining and measuring proximity to criticality

**DOI:** 10.1101/2025.08.03.668332

**Authors:** J. Samuel Sooter, Antonio J. Fontenele, Andrea K. Barreiro, Cheng Ly, Keith B. Hengen, Woodrow L. Shew

## Abstract

Over the half century since the renormalization group (RG) brought about deep understanding of critical phenomena in condensed matter physics, it has been claimed that diverse social, engineered, astrophysical, and biological systems operate close to criticality. However, these systems do not afford the neat phase diagrams and exquisite control available in condensed matter physics. How can one assess proximity to criticality when control parameters are unknown, difficult to manipulate experimentally, and fluctuating in response to changing environmental or internal conditions? Here we meet this challenge with a rigorous theoretical framework and data-analytic strategy for measuring proximity to criticality from observed system dynamics. We developed a temporal RG, well-suited to commonly measured time series, and an information theoretic quantification of proximity to criticality that is independent of model parameterization. After benchmarking our approach on diverse ground-truth cases, we apply it to recordings of spiking activity in the mammalian brain, addressing a long-standing controversy. We show that brain dynamics shift closer to criticality during wakefulness and shift away during deep sleep.

---

Science makes sense of the world by inferring models from data and then extracting meaning from those models. Successful models distill essential features of the data, discarding as many irrelevant details as possible [1–4]. A compelling argument for the feasibility of this program comes from the renormalization group (RG) [5–7]. Originally developed to understand the macroscopic behavior of systems near a phase transition, RG is a theoretical framework for tracking how models change under a change of scale. Surprisingly, RG tells us that most details do not matter; only a few essential model parameters control its macroscopic behavior. This fact suggests that diverse systems - biological, astrophysical, engineered, etc - could share common governing principles despite vastly different details.

From a geometric perspective, RG maps out the space of all models - each point in this space is a model and each dimension is a model parameter (Fig. 1) [8–10]. RG partitions model space into sets of models sharing the same coarse-scale behavior. These sets of models are determined by a mapping on model space that transforms a model into a coarse-grained version of itself. Iterating this mapping creates a trajectory, or flow through model space. Fixed points of this mapping are models that are invariant under coarse-graining, i.e. scale-invariant. Excepting “trivial” fixed points (those without correlated fluctuations), the basins of attraction of fixed points usually coincide with boundaries in model space, where small changes in parameters cause large changes in behavior - i.e. phase diagram boundaries. A model lying in one of these basins is at “criticality.”

**FIG. 1.**
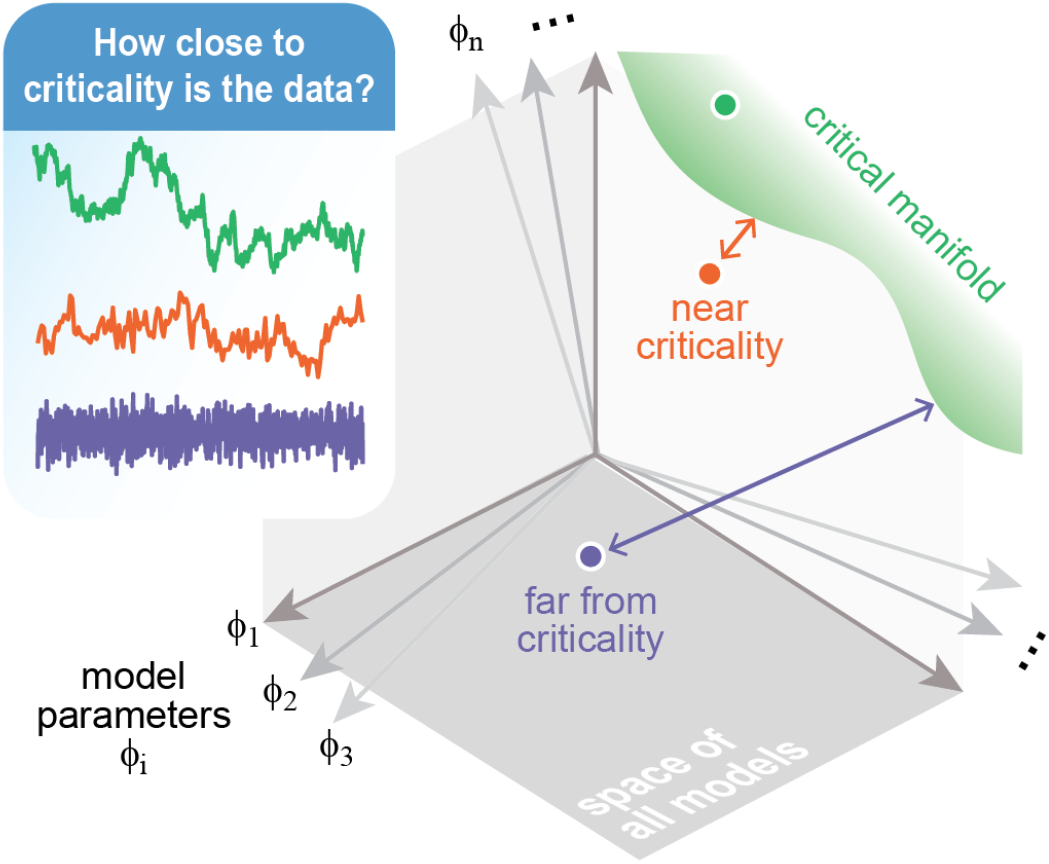
Principled measurement of proximity to criticality. Given a data time series, we aim to assess its proximity to criticality without knowledge of the control parameters (*ϕ*_*i*_) of the system that generated the data. We first develop temporal RG to determine where criticality lies (green) in the space of all models. We then use an information theoretic model comparison (KL divergence rate, double-headed arrows) to quantify how far criticality is from a model that best fits the data (filled circles).

Theoretical predictions abound for how systems behave at criticality: spatial and/or temporal correlations span an infinite range of scales and power-law dependencies emerge with exponents that obey scaling relations [11–13]. Experiments that approximately confirm these predictions have led to claims of criticality in diverse systems from solar flares [14, 15] to social media [16] to flocks of birds [17, 18] to the brain [19–21] and many more. However, these systems have largely unknown phase diagrams and experimentally inaccessible control parameters, which erodes confidence in claims of criticality. This problem is pronounced for biological systems, which self-tune their control parameters in response to their environment. Of particular interest is the hypothesis that the mammalian brain operates near criticality, with important implications for cognition and health [11, 12, 22, 23]. Observations of approximate power-law fluctuations and predicted scaling laws support this hypothesis (systematically reviewed by [23]). The brain self-tunes its state on the fast timescales (seconds-minutes) of neuromodulatory and behavioral state changes [24–28] and on slower timescales (hours-years) of structural plasticity and disease progression [29, 30]. To make stronger claims about criticality in the brain, we require a way to track proximity to criticality without access to control parameters. A zoo of ad hoc tools for detecting criticality in neural systems have been proposed [23]. For example, neuronal avalanches (large fluctuations of collective neural activity) are expected to be power-law distributed at criticality [19, 21] and to obey the “crackling noise” scaling relation [9, 31]. Thus, deviations from power-law distributions [32–36] and/or scaling relations [21, 25, 29, 30, 37, 38] have been quantified and interpreted as deviation from criticality. Other approaches include branching ratio estimates [19, 39–41], measuring the flatness of an effective potential [42], tracking how statistics change upon phenomenological coarse-graining [20, 43–47], and examining system timescales [48–52]. Closest to our approach here, many have fit models to data and examined how best-fit parameters differ from parameters values at criticality [53–56]. However, this approach is unsatisfying because it depends on the arbitrary choice of how to parameterize the model.

Here we propose a parameter-free definition of distance to criticality, rigorously grounded in RG and information theory. We implement our definition for a fundamental class of dynamical models – autoregressive (AR) models – which are well-suited to fitting experimental data. The result is a powerful tool for directly measuring distance to criticality from time series data. We validate this tool on diverse ground-truth cases and apply it to recordings of brain activity, making time-resolved measurements of proximity to criticality. We show that isocortex is closest to criticality while animals are awake and deviates from criticality during sleep.

## I. THEORY

### A. Defining criticality: temporal RG

Traditionally, RG concerns spatial scale-invariance, because interactions are spatially organized (e.g. near neighbor interactions on an Ising lattice). However, spatial RG is limited for the brain, because neural interactions are largely not spatially organized; *<* 5% of the variance in connection strength among mouse brain regions is explained by their spatial separation [57]. Moreover, measurements of brain activity with single-cell resolution are often spatially restricted (usually ~1 mm^3^). These factors limit empirical assessment of spatial scale-invariance. In contrast, typical measurements are temporally unrestricted, recorded for hours or days with millisecond resolution. What is the appropriate RG scheme for a system in which temporal structure is more accessible than spatial structure?

More concretely, consider a time series *x*_1_, *x*_2_, … representing, for example, the activity of a population of neurons in a small patch of cortex. A model for this data is a probability distribution *P* (*x*_1_, *x*_2_, …) for all possible time series. What does it mean for a time series model to be at criticality? To answer this questions, we require an RG program for time series. By direct analogy with Wilson’s momentum space scheme [6], we define a temporal RG (tRG) consisting of the following steps:

1. Switch to frequency domain: Fourier transform 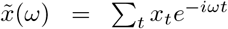 the time series to replace *P* (*x*_1_, *x*_2_, …) with 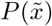.
2. Coarse grain: marginalize over Fourier components with frequency above a cutoff *ω*_0_. Denoting above-cutoff components by 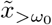 and the remaining low-frequency components by 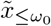, coarse-graining is

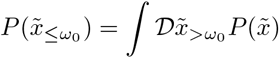

where 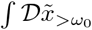 denotes functional integration over all possible 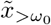.
3. Renormalize: Assessing self-similarity of the original model 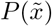 and the coarse-grained one 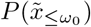 requires rescaling so they have the same range of frequencies,

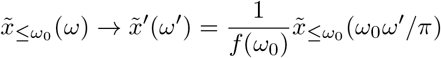

The rescaled function 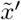 has the same domain of definition, namely [−*π, π*], as the original 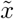. Writing 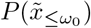 in terms of 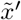, we obtain the fully renormalized distribution 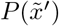.

A typical time series model has a finite autocorrelation time, say *τ*. Coarse graining beyond *τ* (i.e. cutoff frequency *ω*_0_ ≲ 1*/τ* in step 2) will result in a model without temporal correlations - it flows to the white noise fixed point. By definition, a time series model that flows into any fixed point, except the trivial white noise fixed point, it is at criticality (Fig. 2A,B).

**FIG. 2.**
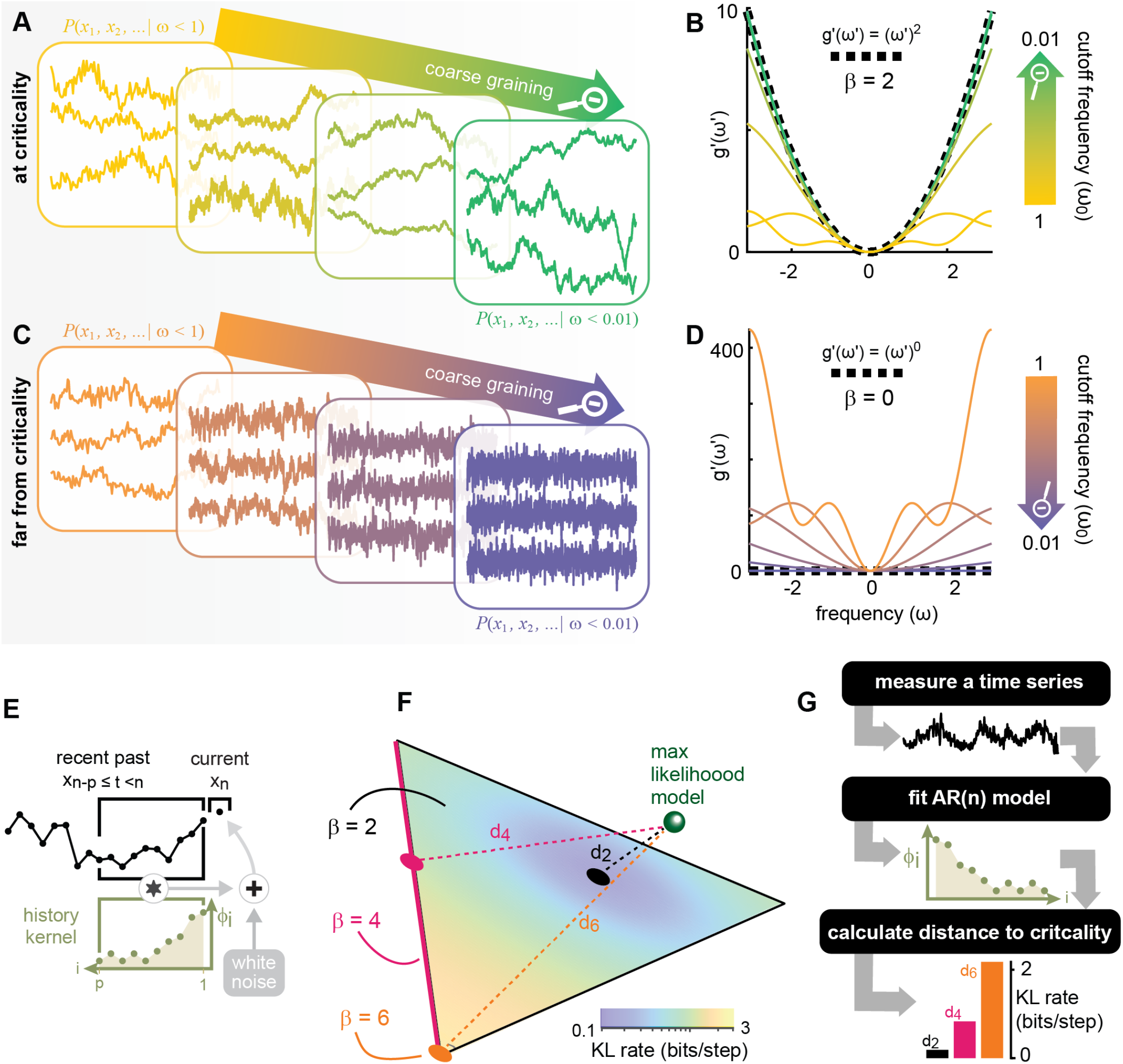
tRG and KL divergence for AR models provides practical tools for measuring proximity to criticality. **(A)** Ensemble of time series generated by a critical AR model (left); applying tRG to this ensemble yields a new ensemble of time series corresponding to a coarse-grained version of the original model (right). **(B)** tRG flow of the AR model in (A). Coarse graining, i.e. taking the cutoff frequency *ω*_0_ → 0, makes *g*′(*ω*′) → (*ω*′)^2^ as the model flows into the *β* = 2 fixed point. **(C)** Same as (A), but for a non-critical AR model; this model flows into the white noise (*β* = 0) fixed point under tRG. **(D)** Same as (B), but for the model in (C). **(E)** Anatomy of an AR model. **(F)** The *β* = 2 critical manifold for AR(n) models lies in a patch of an n-1 dimensional hyperplane (in this example, n=3 and the patch is triangular). Along one edge of the *β* = 2 manifold is the (*n* −2)-dimensional *β* = 4 manifold, along one edge of which is the (*n* − 3)-dimensional *β* = 6 manifold, and so on. Distances to criticality - *d*_2_, *d*_4_, and *d*_6_ - are sketched for an example model (green point). The nearest critical models for each type of criticality are the colored dots on the critical manifold. The surface color of the *β* = 2 manifold indicates the KL distance from the example model to each point on the *β* = 2 manifold. **(G)** Data analytic pipeline: step 1 - acquire some time series data, step 2 - obtain max likelihood fit AR(n) model, step 3 - use KL rate and our tRG for AR models to measure distances to each tRG fixed point.

### B. Defining distance to criticality: KL divergence

Given a time series model, *how close* is it to criticality? More specifically, if one model represents some data and another model is at criticality, how does one define distance between the two models in a way that does not depend on an arbitrary choice of model parameterization? Information theory supplies a natural choice; Kullback-Leibler (KL) divergence [58, 59] per unit time, defined for two models *P*_A_(*x*_1_, *x*_2_, …) and *P*_B_(*x*_1_, *x*_2_, …) as

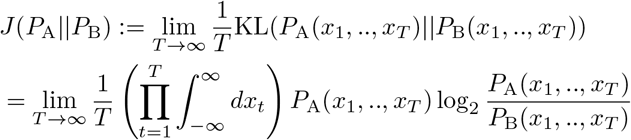

where *P*_A_(*x*_1_, …, *x*_*T*_) denotes the full model *P*_A_(*x*_1_, *x*_2_, …) marginalized over all *x*_*t*_ with *t > T*, and similarly for B.

We now *define* the distance to criticality of a time series model A as

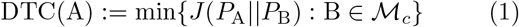

where ℳ_*c*_ is the set of all time series models at criticality (i.e. the set of all models that flow into a non-trivial tRG fixed point). This is the distance between model A and the nearest critical model, where “nearest” means minimal KL divergence rate. This quantity has units of bits per time and can be interpreted as how distinguishable (per unit time) model A is from the nearest critical model. More explicitly, DTC(A) is the expected rate of accumulation of evidence for the hypothesis “the underlying model is A” over the alternative hypothesis “the underlying model is at criticality” when observing a time series generated by A. This definition of DTC is parameter-free (i.e., it depends on the model dynamics, not the model parameters) and has a physically meaningful interpretation. However, working with the set of all critical models ℳ_*c*_ is infeasible; in practice, we replace ℳ_*c*_ with its intersection with a tractable family of models ℳ (see Supp. Mat. S6 for further discussion). For example, in Section I.C we carry out this program analytically with ℳ equal to the family of all autoregressive models of a given order. Section II employs our DTC definition in other families of models, including ones that incorporate spiking, nonlinearity, and multiplicative noise.

### C. tRG and DTC for AR models

Measuring DTC from time series data presents two main challenges: (i) inferring an unknown model underlying the data and (ii) determining the structure of the set ℳ_*c*_ of critical models. A celebrated approach to challenge (i) is maximum entropy modeling (*max ent*) [60]. *Max ent* acknowledges that, given limited samples, only low-order statistics, like means and covariances, can be measured accurately. Thus, we seek the minimally presumptive model consistent with these low-order statistics. For time series, the *max ent* model that reproduces measured autocovariances *Ĉ*(*t*) for lags *t* = 0, 1, …, *n* is the order-*n* AR model (Fig. 2E)[61]:

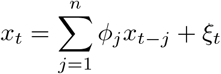

where the history kernel *ϕ* = (*ϕ*_1_, …, *ϕ*_*n*_)^*T*^ is given by

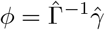

and *ξ*_*t*_ is Gaussian white noise with variance

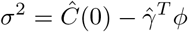

Here 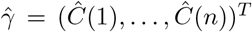 and 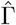 is an *n × n* matrix with 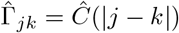.

In systems with uncontrolled control parameters, like the brain, proximity to criticality is time-dependent. To track such changes, we must measure proximity to criticality in many consecutive short duration periods (e.g. in a sliding window). Considering how few samples are in each short window, AR models are fundamental to this endeavor (in the *max ent* sense). We next perform a full tRG analysis of AR(n) models, identifying all the fixed points and their basins of attraction. (RG for AR(2) was previously developed with a different approach [10].) Moreover, we analytically calculate the KL divergence rate between any two AR models, allowing us to fully realize our rigorous definition of distance to criticality.

The probability distribution corresponding to an AR model with history kernel *ϕ* and noise variance *σ*^2^ is (Supp. Mat. S2.1)

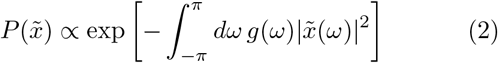

where

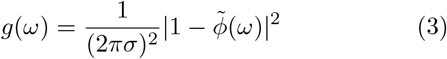

This makes tRG coarse-graining straightforward; the high and low frequency components factor separately, leaving us with

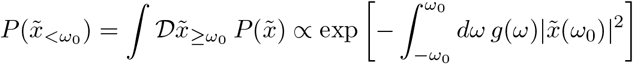

After rescaling, this becomes

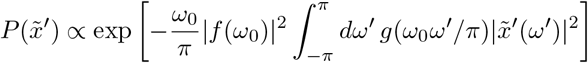

which is the same as the original distribution 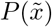 with *g*(*ω*) replaced by

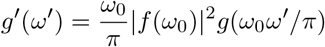

As *ω*_0_ → 0^+^ (i.e., coarse-graining more and more), the behavior of *g*′(*ω*′) is determined by the dominant term in its Taylor series expansion, which is an even power of *ω* because *g*(*ω*) is an even function of 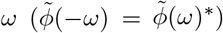. Thus, we can write *g*(*ω*) = *Aω*^*β*^ + 𝒪 (*ω*^*β*+2^), where *β ∈* {0, 2, 4, …}. Then choosing *f* (*ω*_0_) = *A*^−1*/*2^(*ω*_0_*/π*)^−(*β*+1)*/*2^ makes

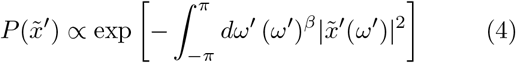

in the *ω*_0_ → 0^+^ limit. We conclude that each value of *β* corresponds to a different tRG fixed point, and the exponent of the dominant term in *g*(*ω*) determines which fixed point an AR model will flow into. Fig. 2B,D shows how *g*(*ω*) changes upon coarse-graining for two example fixed points.

Determining the sets of AR models that flow into each fixed point then reduces to identifying conditions for the dominant term in *g*(*ω*) to have a given exponent. We show in Supplementary Materials (S2.2) that an AR model flows into the *β* fixed point (Eq. (4)) for *β ≥* 2 if and only if its history kernel *ϕ* satisfies (i) *a*_0_ = 1, (ii) *a*_*m*_ = 0 for all *m ∈* {1, 2, …, *β/*2 − 1}, and (iii) *a*_*β/*2_ ≠ 0, where

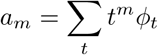

If instead *a*_0_ ≠ 1, then the model flows into the trivial *β* = 0 fixed point - the white noise fixed point. The remaining fixed points are non-trivial; models that flow into them exhibit fluctuations across all timescales. In RG parlance, the parameters *a*_0_, *a*_1_, …, *a*_*β/*2−1_ are *relevant* for the *β* fixed point. Changing any other details of an AR model, including its noise variance, does not impact its asymptotic behavior at coarse scales. This results in extended basins of attraction of the tRG fixed points in the space of possible history kernels *ϕ* (Fig. 2F).

Now consider a specific AR model, say A. How far is A from criticality? The KL divergence rate between A and another AR model B is (Supp. Mat. S2.3)

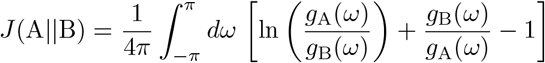

where *g*_A_(*ω*) and *g*_B_(*ω*) are defined as in Eq. (3). To calculate A’s distance to criticality, we numerically minimize *J*(A || B) over all critical AR models B of the same order as A (Fig. 2F; Supp. Mat. S2.4). We can also calculate A’s distance to each *type* of criticality that AR models exhibit; specifically, for each *β*, we define *d*_*β*_ as the minimum of *J*(A || B) over B in the set of critical AR models of type *β*. The nested structure of the tRG basins guarantees that *d*_2_ ≤ *d*_4_ ≤ *d*_6_ ….

With this practical implementation of our definition of proximity to criticality, we arrive at a two-step pipeline for measurements of distance to criticality for real data (Fig. 2G). Step 1: fit an AR model to the real data. Step 2: use our tRG results and formula for KL divergence rate to measure the distance from the best fit model to the nearest model on the critical manifold.

## II. BENCHMARKING

While AR models are justified in the *max ent* sense, it is not immediately obvious that AR models are good fit for brain data. Neurons are nonlinear, population activity is sometimes highly non-Gaussian, and many important experimental measurements are of spikes. In contrast, AR models are linear, Gaussian, and do not directly model spikes. Can our approach handle data that is not perfectly fit by an AR model? Next we show that our tools work surprisingly well in three cases: 1) a point process model that generates spike-like data (Hawkes process, Fig. 3A-C), 2) a nonlinear time series model (overdamped Langevin equation, Fig. 3D,E), and 3) a non-Gaussian bursty model (Fig. 3F,G). For these models, critical parameter configurations are known (Hawkes: *A* = 1, *µ* = any; Nonlinear: *b* = *c* = 0; Bursty: *b* = 0, *h* = any). We compared *d*_2_ to the ground truth DTC for each model, which we obtained by analytical and numerical methods (Supp. Mat. S3.1-S3.4). True DTC was faithfully revealed by *d*_2_ in all cases except for very bursty dynamics (red dashed area Fig. 3G). We conclude that AR models work well for our goals, but not without limitations.

**FIG. 3.**
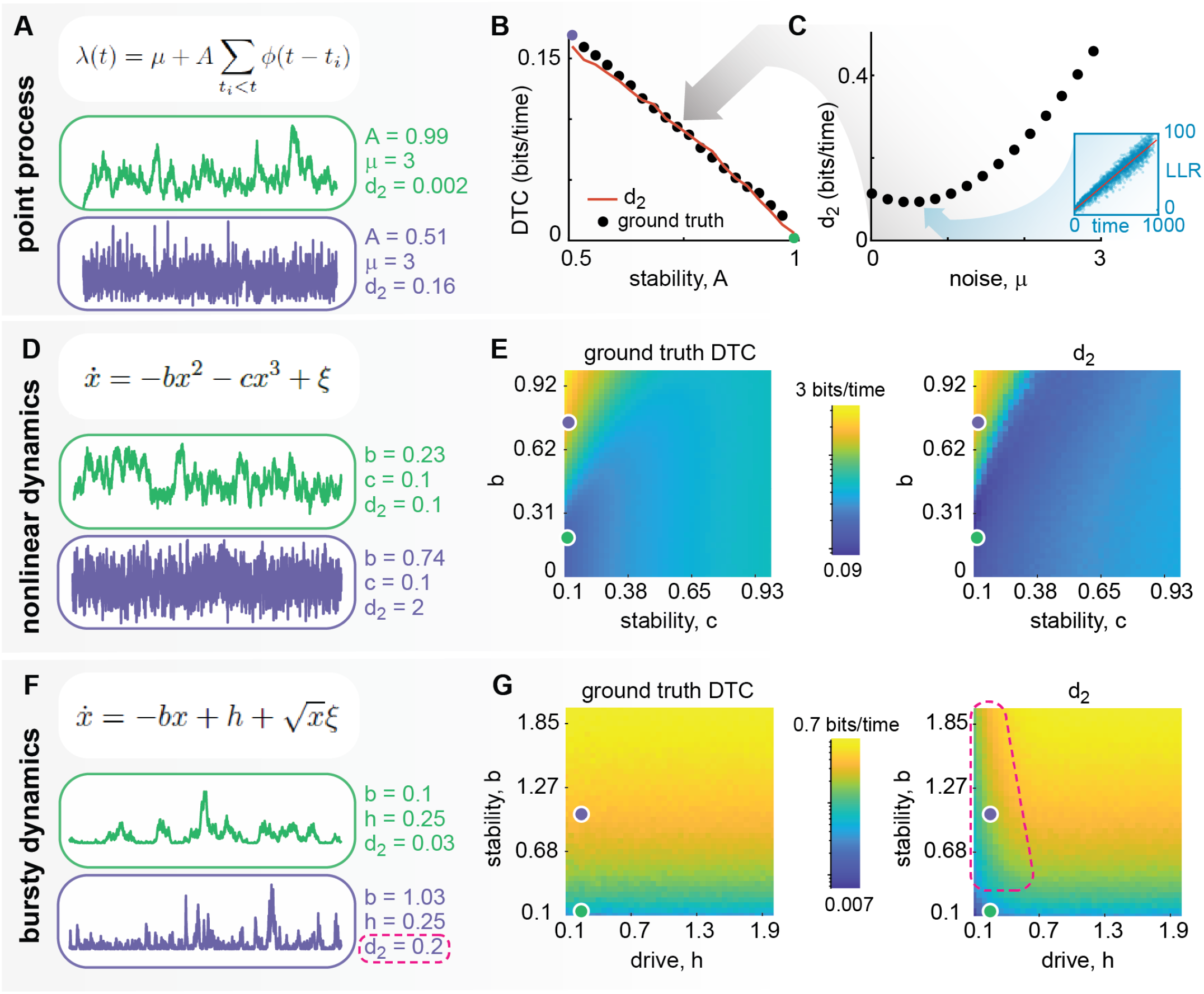
Benchmarking *d*_2_ against ground truth cases. **(A)** A Hawkes process is at criticality for *A* = 1, for all *µ*. Example “spike” count time series near criticality (green) and far from criticality (purple). **(B)** *d*_2_ closely matches ground truth DTC across a wide swath of parameter space, including models far from criticality (*A* ≪ 1) and models close to criticality (*A ~* 1). **(C)** The ground-truth DTC values in (B) were calculated by minimizing the KL divergence rate over the set of critical Hawkes processes. Inset: numerical calculation of KL divergence rate. **(D)** Langevin dynamics in a quartic potential is at criticality for *b* = *c* = 0. **(E)** Ground truth DTC agrees well with *d*_2_ as a function of *b* and *c*. **(F)** The bursty model is at criticality for *b* = 0, for all *h*. **(G)** *d*_2_ matches ground truth DTC well, except when *h* is small (red dashed line), which is when the model dynamics deviate extremely from Gaussian, precluding a good AR model fit.

## III. APPLICATION TO CORTICAL SPIKING ACTIVITY

As a first use case for our approach, we asked whether the mammalian brain is closer to criticality during sleep or during wake. Previous studies report inconsistent results; some suggest sleep is closer to criticality than wake [62], others suggest the opposite [51, 52], and some report little difference [27, 63–65]. Here, we performed electrophysiological recordings of single-unit spike activity from primary visual cortex of freely-behaving rats (first reported in [27]) and analyzed equivalent data from mice (first reported in [66]). We analyzed 96-240 h of data recorded in each of N = 8 rats, with a mean of 35 ± 7.9 single units per 12 h clustering block (see [27] for details). For the mice, we analyzed 13 recordings, 6.2 ± 1.8 hours in duration (mean ± sd), 67 ± 18 neurons per recording. In both datasets, periods when the animals were awake, in deep sleep (NREM sleep), or in REM sleep were determined based on animal movement and/or local field potential measurements as detailed in the original papers [27, 66]. We measured proximity to criticality (*d*_2_) during different states of sleep and wake. First, we created a spike count time series (Δ*t* = 40 ms time bins). We fit an AR model to this spike count time series separately for awake (Fig. 4A), NREM sleep (Fig. 4B), and REM sleep periods (Fig. 4C) (Methods). For both mice and rats, cortical neurons were closest to criticality (*d*_2_ was smallest) in the awake state and furthest from criticality during NREM sleep (Fig. 4D). Considering other tRG fixed points, *d*_*β*_’s with *β* = 4, 6,.. also distinguish the sleep and wake states (Supp. Fig. S1).

**FIG. 4.**
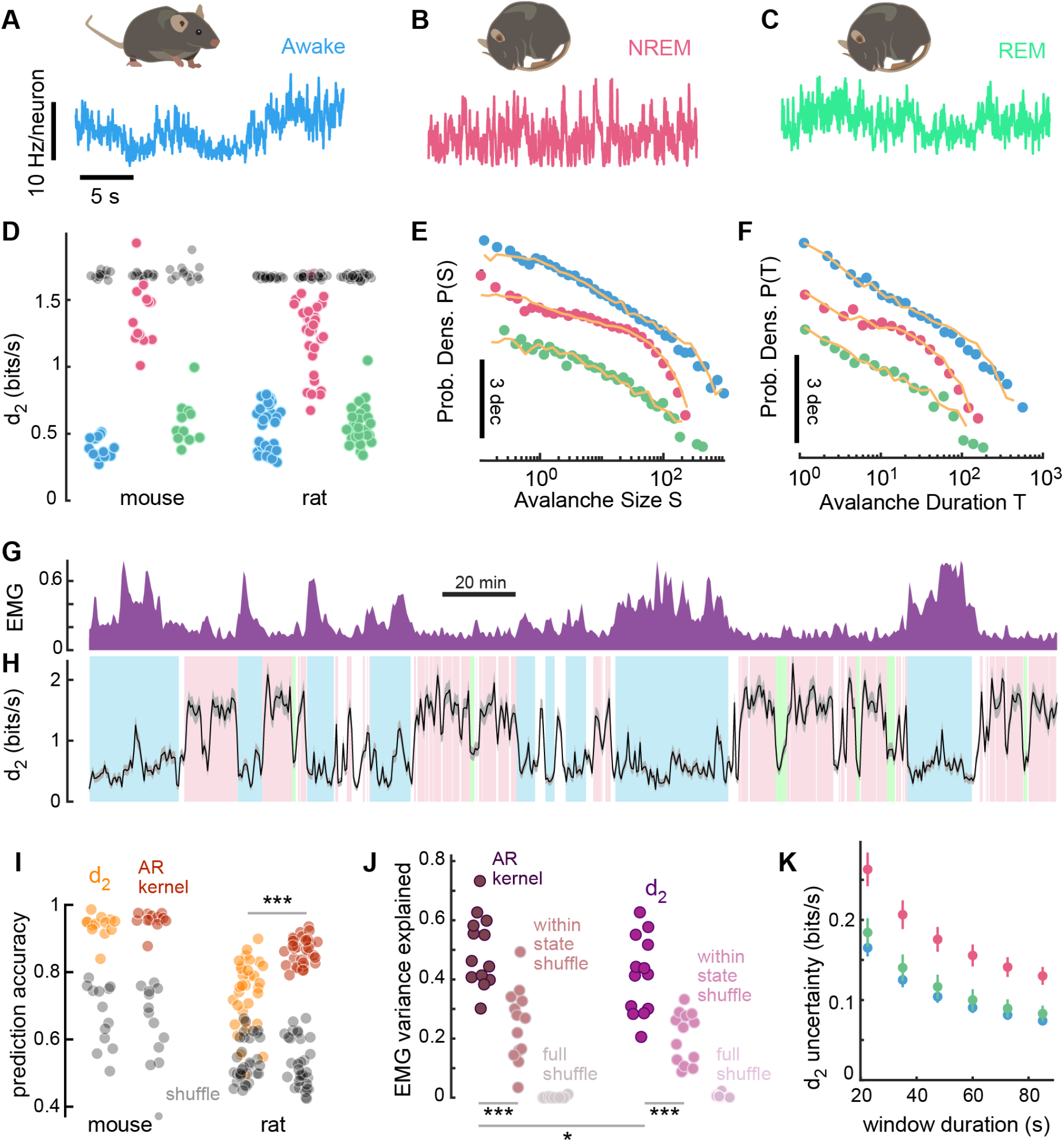
Proximity to criticality in mammalian brain depends on vigilance state and behavior. **(A)**-**(C)** Mouse V1 activity during wakefulness, NREM sleep, and REM sleep. Each row of dots represents spike times from a single neuron. We apply our approach to spike count time series (colored lines). **(D)** V1 is closest to criticality (*d*_2_) during wake (blue) and furthest during NREM (red). Faded gray dots are for rate-matched Poisson surrogate data. Each point represents a single recording session. **(E)** Avalanche size distributions for mouse V1 during each behavioral state (markers) match well to avalanche size distributions from simulations of best-fit AR models (yellow lines). These examples are from the same recording session as shown in (A)-(C). **(F)** Same as (E), but for avalanche duration. **(G)** Body activity (electromyography, EMG), obtained simultaneously with the *d*_2_ timeseries in (H). **(H)** *d*_2_ calculated in a 60 s sliding window reveals time-resolved shifts in proximity to criticality (black line). Gray shaded area represents uncertainty Δ*d*_2_. Color of background indicates wake (blue), NREM (red), or REM (green) periods. **(I)** Sleep/wake states can be predicted with high accuracy using *d*_2_ (orange) and even higher accuracy using all AR model coefficients (red). **(J)** Body movement (EMG) can be predicted using *d*_2_ nearly as well as prediction based all AR model coefficients (\**p <* 0.05). EMG fluctuations within each sleep/wake state are predicted by *d*_2_ (\*\*\**p <* 0.001), as evidenced by drop in prediction accuracy when EMG is shuffled within each state. **(K)** Uncertainty in *d*_2_ depends on sleep state and decreases for increasing duration of the sliding window duration.

For more direct comparison to previous studies, we also performed avalanche analysis (Methods). We created distributions of avalanche sizes and durations and used them for two purposes. First, in agreement with minimal *d*_2_ in the wake state, we found that avalanches had the largest range of power-law scaling in the awake state (Fig. 4E,F, Supp. Fig. S2-S4 for exhaustive results). Second, we used avalanche distributions as an additional check of how well the fitted AR models matched the observed dynamics. Close agreement between the avalanche distributions of the experimental data and those obtained from simulations of the best fit AR models (yellow lines in Fig. 4E,F) enhances confidence in our approach.

These results summarize multi-hour recordings. Can our approach track changes in proximity to criticality in a time-resolved way? We performed *d*_*β*_ measurements in a sliding 60 s window and found that *d*_2_ fluctuations reflected changes in sleep/wake states and body movement (EMG) (Fig. 4G,H). The uncertainty in *d*_2_ (Supp. Mat. S2.5) is small compared to *d*_2_ changes across sleep/wake states for 60 s or even shorter sliding window durations (Fig. 1J). We could predict sleep/wake states with 94 ± 1% and 74 ± 2% accuracy in the mouse and rat data, respectively (mean ± sem across sessions, Methods). More impressive, we were able to predict EMG using *d*_2_, including EMG fluctuations within each sleep/wake state (Fig. 4I). Overall, these results demonstrate that *d*_2_ can track proximity to criticality, even in living systems with slewing underlying control parameters.

## IV. DISCUSSION

We have proposed a principled definition of distance to criticality and implemented it by developing a temporal RG and KL divergence rate for AR(n) models. We leveraged this theoretical framework for measuring distance to criticality to address a long-standing controversy, showing that cortex is closer to criticality during wake than NREM sleep in two independently collected datasets from two different species (mice and rats).

For a given time series, the repertoire of timescales present in the fluctuations depends strongly on proximity to criticality. Thus, *d*_2_ depends strongly on the timescale repertoire. How does *d*_2_ compare to other assessments of the timescale repertoire? These include power spectra [67, 68], dominant timescale estimates [41, 51, 52], detrended fluctuation analysis (DFA) [48, 49], avalanche analysis [19, 21] and more [40]. A comprehensive quantitative comparison will require further work, but we mention a few important differences. First, avalanche analysis focuses on statistics of large fluctuations - the avalanches - and is blind to what is happening between avalanches - the “quiet times”. In contrast *d*_2_ accounts for the entire timeseries, thus accounting for temporal scale invariance differently than avalanche analysis. This could explain why some avalanche-based analyses did not find differences between wake and NREM sleep [27], while *d*_2_ reveals a difference (Fig. 4D,H). An advantage over avalanche analysis is that *d*_2_ works on short time series, affording a time-resolved, sliding-window assessment of proximity to criticality (Fig. 4G). Comparing to methods that probe the dominant timescale of a time series [41, 51, 52], a decrease in *d*_2_ is likely when the dominant timescale increases, as expected for “critical slowing down”. However, distance to criticality is not determined solely by the longest timescales of the system; for a fixed longest timescale, *d*_2_ can vary over an order of magnitude (Supp. Fig. S6). Comparing to power spectra and DFA, we note that their power-law exponents are related to criticality, but do not measure proximity to criticality. This is particularly true when the shape of the power spectrum or fluctuation function deviates from a powerlaw, which is expected for a system that departs from criticality.

There are many open questions about proximity to criticality and brain function. How do deviations from criticality relate to brain disorders [23, 30, 69, 70]? Does distance to criticality vary across brain regions [71, 72]? Do moment-to-moment shifts in proximity to criticality impact cognitive performance [52]? We have established a solid theoretical foundation for these questions and theory-driven data analytic tools (and freely available code: *github*…*doi*…*TBD*) to answer them using experimental data.

## V. METHODS

### Rat experiments

Rats were implanted with an 64-electrode array (16 tetrodes made of 12 *µ*m wires) in primary visual cortex under isoflurane anesthesia as detailed previously [27]. Implantation coordinates were 1.45 mm/3.45 mm/ −1 mm (anteroposterior/mediolateral/dorsoventral relative to lambda and dura). The total headstage weight, including cement, bone screw andenclosure, was 2 g. After 2 days of recovery, neural activity was recorded (25 kS/s, eCube Server, White Matter) continuously for 10–14 days during free behavior in home cage. Animal behavior was monitored with 15 or 30 fps video recording synchronized to the electrophysiological data with Watchtower software (White Matter). Raw electrophysiological data were bandpass filtered between 500 and 7500 Hz and thresholded for spike-waveform extraction (mean ± 4 s.d.). Spike sorting was performed with a modified version of SpikeInterface63 and MountainSort4 as described in [27].

### Mouse experiments

As reported previously [66], mice were implanted with a 64-channel silicon probe (single shank, Cambridge NeuroTech, H3 64x1 probe) mounted on a movable microdrive for recording the activity of multiple single-units and local field potentials (LFPs) in primary visual cortex. The probe was implanted at 1.0 mm/2.5 mm anteroposterior/mediolateral, with a 21 degree angle from dorso-ventral axis and a 10 degree angle from the mediolateral axis. The probe tip was 1.0 mm deep after insertion. After overnight recovery in their home cage, extracellular electrophysiological recording were performed while the animals slept or walked around freely moving in the home cage for 68 hr. Electrophysiological data were acquired using an Intan RHD2000 system (Intan Technologies LLC) sampled at 20 kHz rate. Spike sorting was performed semi-automatically, using Kilosort.

### Calculating d_2_ from segmented time series

In Fig. 4D, we reported a single *d*_2_ value for each state (wake, NREM, or REM). To obtain a *d*_2_ for NREM, for instance, we first calculated the autocorrelation function separately for each NREM segment and then averaged across all segments (accounting for variable lengths of segments). Then we used the Yule-Walker method to obtain best-fit AR model coefficients from the average autocorrelation function. We then used these coefficients to calculate *d*_2_. In contrast, in the time-resolved *d*_2_ analysis in Fig. 4H, we used maximum likelihood estimation (MLE) of the AR model coefficients because this allowed us to estimate error bars for *d*_2_ (Supp. Mat. S2.3).

### Avalanche analysis

To generate the avalanche distributions in Fig. 4E,F and in Supp. Fig. S2–4, we used standard procedures, as described elsewhere (e.g., [21]). In brief, the first step is to create a spike count time series *x*_1_, *x*_2_, …, where *x*_*i*_ is the total number of spikes from all recorded neurons in the *i*th time bin. Time bins are non-overlapping, consecutive, and 40 ms in duration. Then, avalanches are defined as periods of time when the spike count exceeds a threshold (10th quantile). For each avalanche, its size is defined as the area between the threshold and the spike count time series and its duration is defined as the time spent above the threshold. Each recording had 1000s of avalanches. Each size or duration probability density distribution was created using 10 logarithmically spaced bins per decade.

### Predicting sleep states

We used a multinomial logistic regression (Matlab’s mnrfit function) to predict sleep state from *d*_2_ and the AR history kernel (Fig. 4I). The model linearly combines the predictor variables (either *d*_2_ or the kernel) to compute a score for each class (NREM, REM, wake), converts these scores into class probabilities using the softmax function, and outputs the class with the highest probability. We evaluated model performance with 5-fold cross-validated classification accuracy.

### Predicting EMG, gradient-boosted decision trees

We used gradient-boosted decision trees (XGBoost algorithm) to predict body movement (electromyogram, EMG; Fig. 4J) from *d*_2_ and the AR history kernel. To select hyperparameters without data leakage, we used nested, temporally contiguous cross-validation. Specifically, we first split each session into temporal halves (“outer” cross-validation folds). We then split each half into two further temporal halves (“inner” folds). Next, we used Bayesian optimization (BayesSearchCV function from scikit-optimize package) to select hyperparameters that maximized cross-validated performance (measured as the mean squared prediction error) over the inner folds. Finally, we reported the model performance as the cross-validated performance across the outer folds. We selected hyperparameters from the following ranges: ‘learning rate’ (0.01, 0.3), ‘n estimators’ (50, 500), ‘max depth’ (3, 10), ‘subsample’ (0.5, 1), ‘colsample bytree’ (0.5, 1), ‘reg alpha’ (0, 10), ‘reg lambda’ (0, 10).

## SUPPLEMENTARY MATERIALS

## 1. Additional details of mouse and rat spike data analysis

### 1.1. Distances to other fixed points - *β >* 2

In the main text, we reported *d*_2_ measurements for the spike data in different sleep and wake conditions. We focused on *d*_2_, because it is the most general distance to criticality, but our tools also measure distances to other tRG fixed points *β* = 4, 6, … up to *β* = 2*n*, where n is model order. In Supplementary Fig. 1, we show these additional *d*_*β*_ measurements. Notice that differences between the three sleep/wake states are apparent not just for *d*_2_. For instance, *d*_6_ seems to distinguish wake versus REM better than *d*_2_. The gray rectangles indicate that our algorithm for computing distance to the nearest (min KL divergence rate) critical model did not converge for *β* =16 and 18. This limitation has, in our experience, only impacted these very large *β* cases.

### 1.2. Avalanche distributions

For comparison to previous studies based on avalanche analysis, we performed avalanche analysis on the mouse (Sup. Fig. 2) and rat data (Sup. Figs. 3,4). We observe that in the awake state, the avalanche distributions tended to have a greater range of power-law scaling, which is consistent with our findings that wake state is closest to criticality, based on *d*_2_. Furthermore, for each recording, we ran a simulation of the best fit AR model and performed avalanche analysis of the simulated time series. The good match between the simulated and experiment avalanche statistics verifies that AR models are good fits to our data.

## 2. Distance to Criticality for AR Models

### 2.1. Probability Distribution of Trajectories

Fix a stationary AR model with kernel *ϕ* = (*ϕ*_1_, …, *ϕ*_*n*_) and noise standard deviation *σ*. Run the model for *T* time steps to generate a trajectory **x** = (*x*_1_, …, *x*_*T*_), and let *C* be the *T × T* covariance matrix for this process. Then

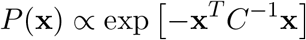

Since this model is stationary, 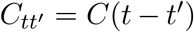. In the large *T* limit, *C* is diagonalized by a Fourier transform. To see this, let *F* be the *T × T* matrix with entries 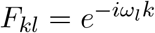, where 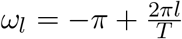. Then

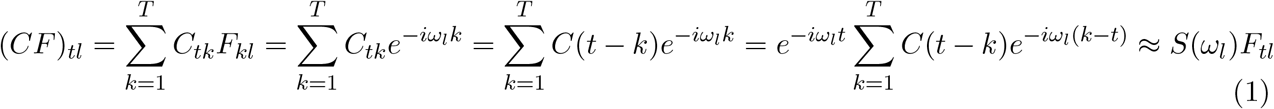

where

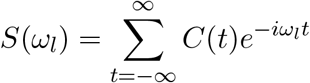

is the power spectral density of the model and in the last step we assumed that *C*(*t*) decays to zero at large lags. In words, Eq. (1) says that Fourier modes are (approximately, for large *T*) eigenvectors of the covariance matrix with eigenvalues given by the power spectral density. But then

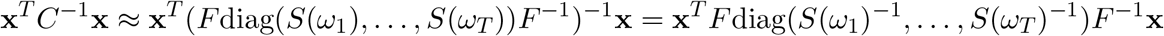

where diag(*S*(*ω*_1_), …, *S*(*ω*_*T*_)) denotes the diagonal matrix with *S*(*ω*_1_), …, *S*(*ω*_*T*_) along the diagonal. Then defining 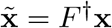 and noticing that 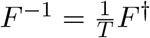, we have

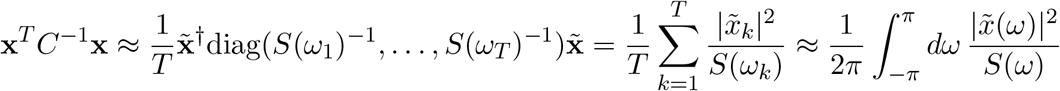

where we defined 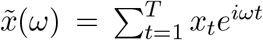. Finally, the power spectral density of an autoregressive process is (see [1], Chapter 4)

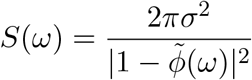

Where 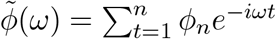, so we arrive at (Eqs. (2–3) in the main text)

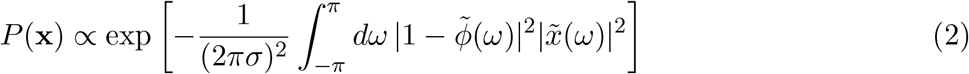

**Figure 1.**
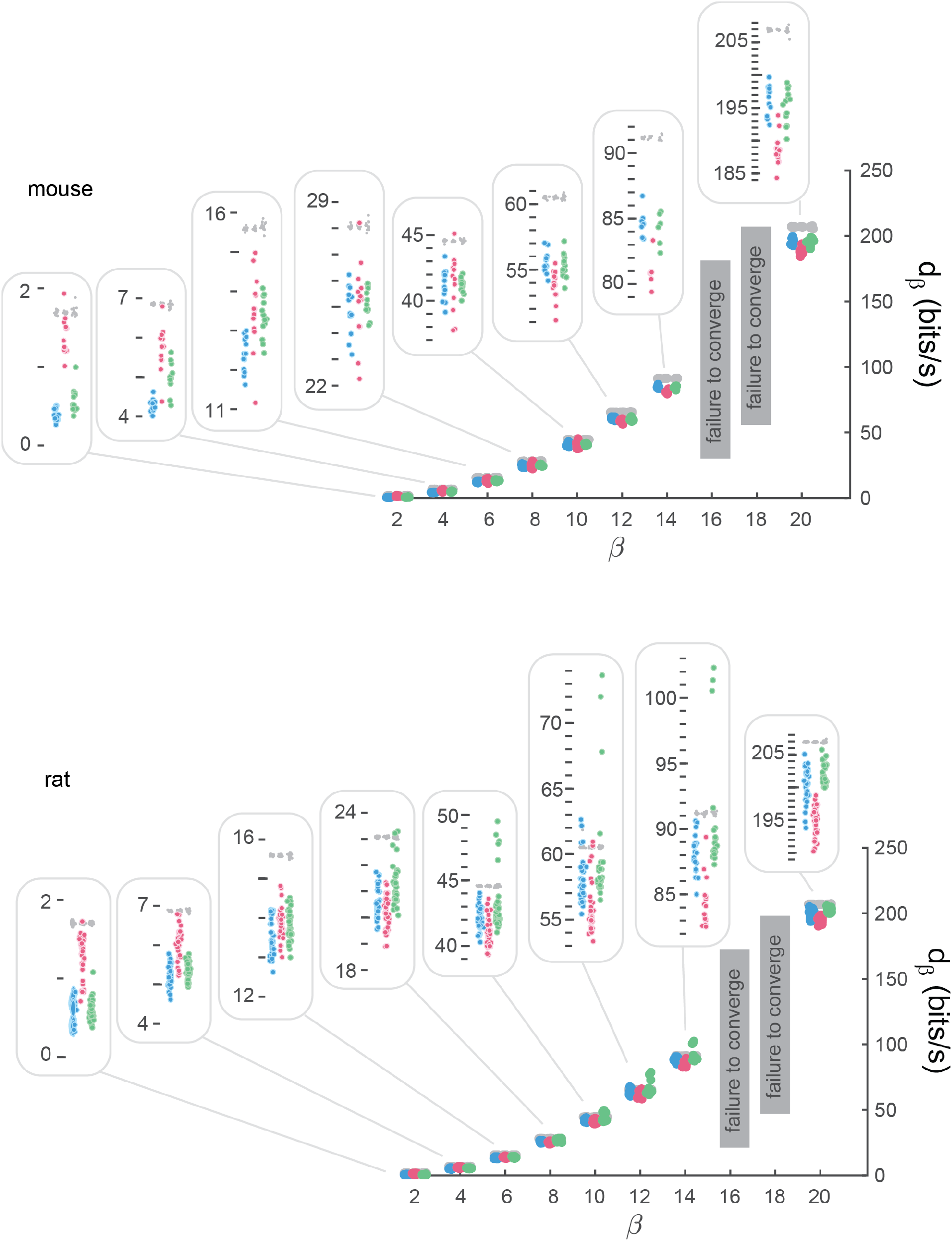
Distances to all fixed points. An AR(10) model was fit to each of the wake (blue), NREM (red), and REM (green) states presented in Fig. 4 of the main text. Thus, there are 10 tRG non-trivial fixed points to consider. In general, distances *d*_*β*_ increase with *β*. Each inset zooms in on one set of *d*_*β*_ results to show sleep versus wake differences.

**Figure 2.**
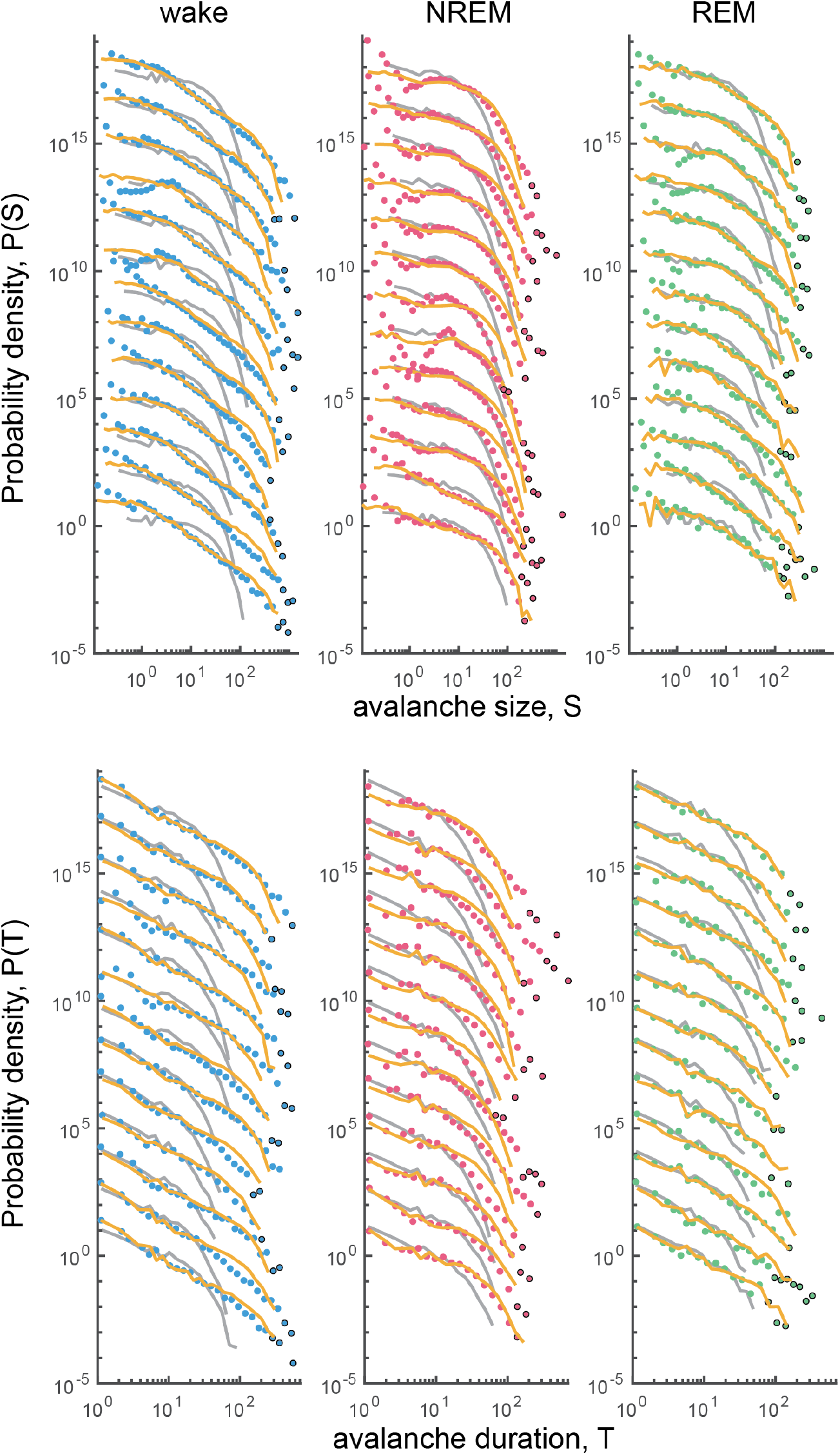
Avalanche distributions for mouse data. **(top left)** Wake state avalanche size distributions for all mouse recordings, offset vertically for comparison of distribution shape. The points represent the original experimental data. Gray lines represent control analyses based on rate-matched Poisson surrogate data. Yellow lines show avalanche distributions from simulations of the best fit AR model. Black circles mark bins containing *≤* 5 avalanches. The match between the yellow lines and the points shows that our AR models are good fit to the data. **(top middle)** Same as (top left), but for NREM periods. **(top right)** Same as (top left), but for REM periods. **(Bottom row)** Same as (top), but for avalanche durations.

**Figure 3.**
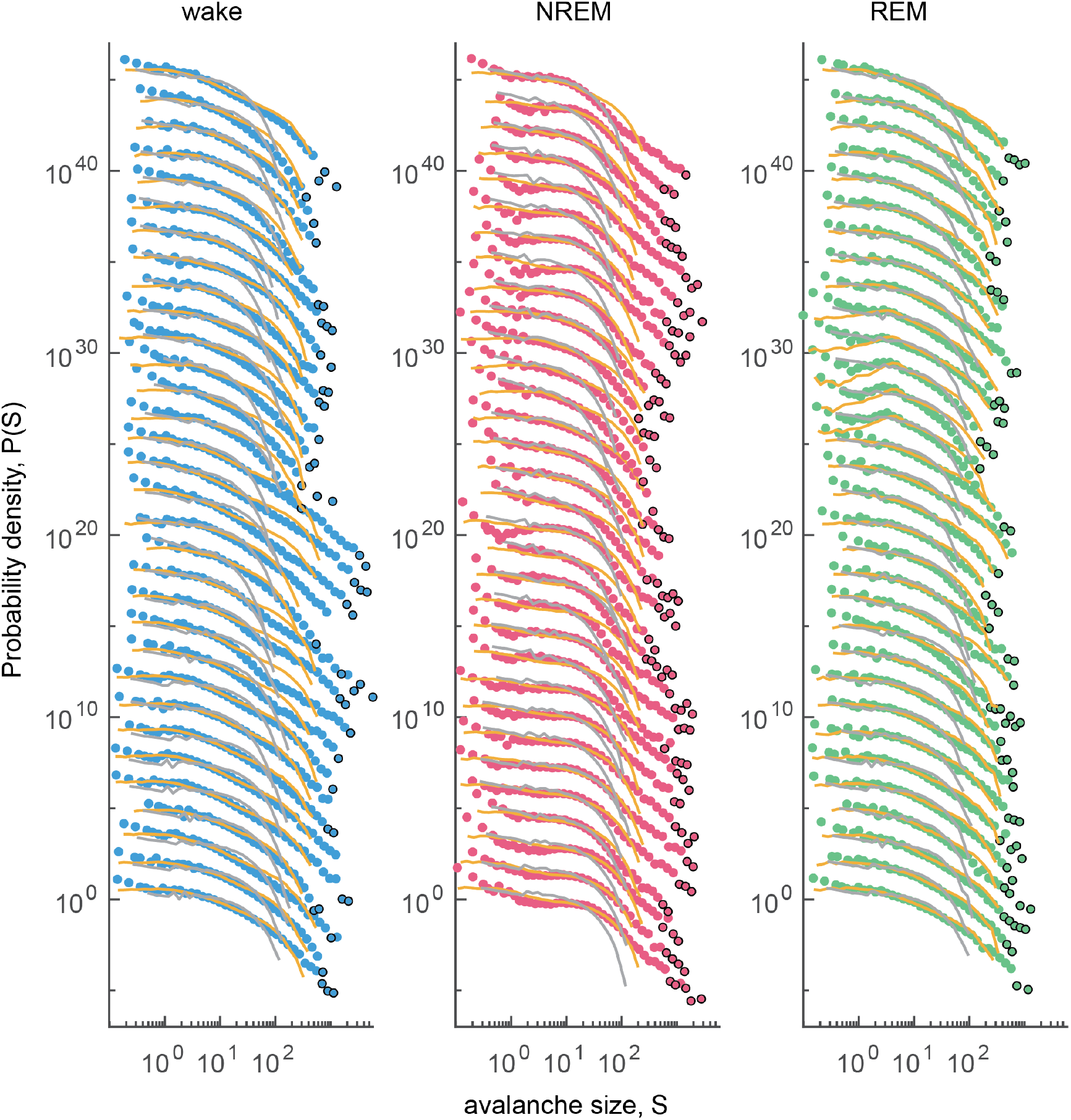
Avalanche size distributions for rat data. Same as top row of Fig. 2, but for rat data.

### 2.2. tRG Basins of Attraction

Fix an AR model (*ϕ, σ*). In Section I.C, we showed that this model flows into the *β* tRG fixed point if and only if *ω*^*β*^ is the dominant term in the Taylor expansion of

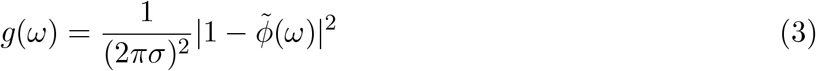

around *ω* = 0. Here we derive the geometry of the sets of AR models that flow into each tRG fixed point. In other words, we locate the basins of attraction of each fixed point in the space of all AR models.

**Figure 4.**
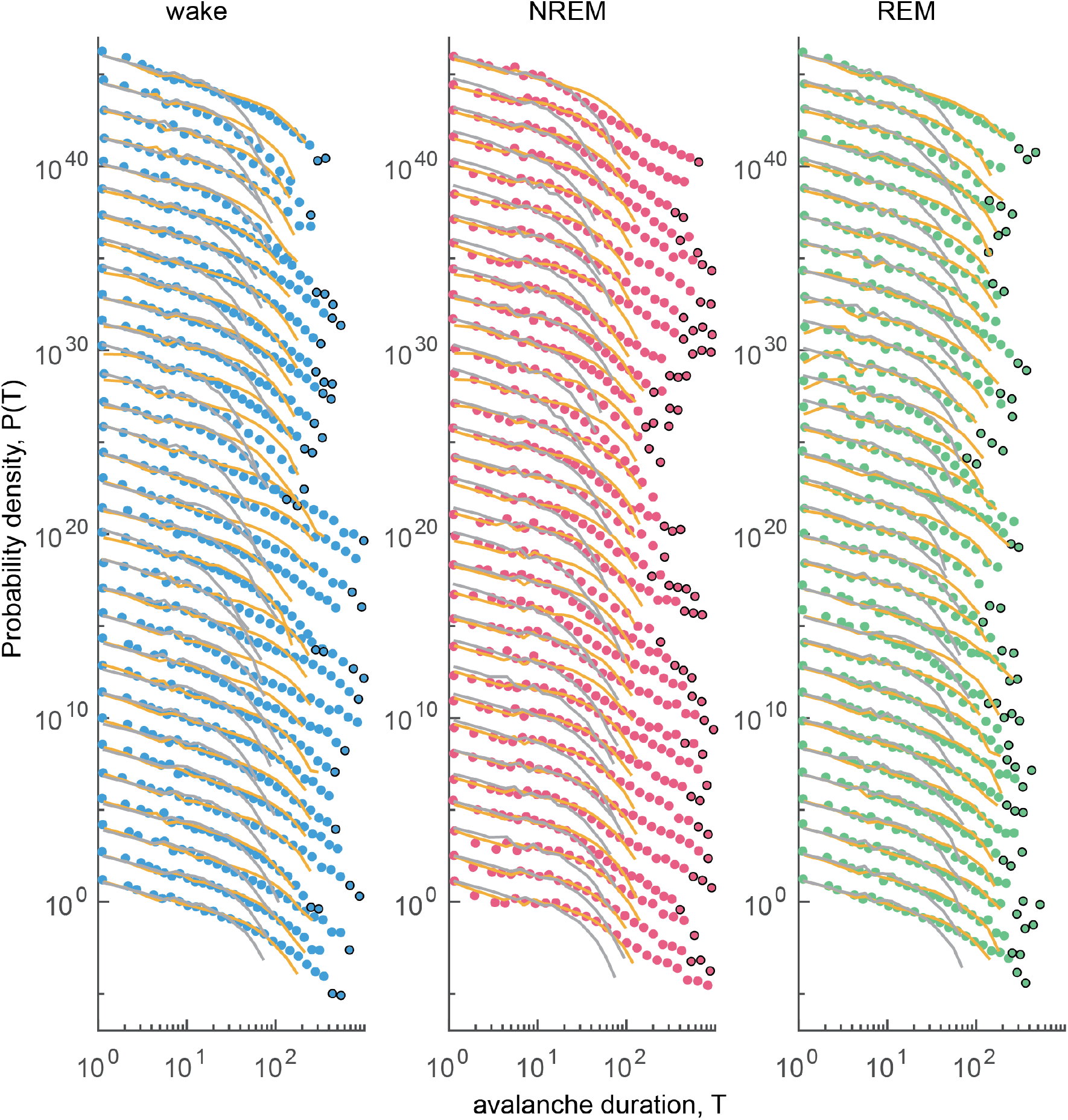
Avalanche duration distributions for rat data. Same as bottom row of Fig. 2, but for rat data.

Let the Taylor expansion of 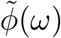 around *ω* = 0 be 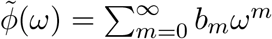. Then

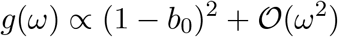

If *b*_0_ ≠ 1, then the dominant term in *g*(*ω*) is the constant term and the AR model flows into the trivial *β* = 0 fixed point. If instead *b*_0_ = 1, then

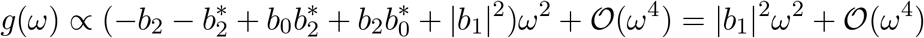

If *b*_1_ ≠ 0, then the dominant term in *g*(*ω*) is the quadratic term, so the AR model flows into the *β* = 2 fixed point. This is the base case of an induction argument for the following proposition:

#### Proposition S2.2.1

For all even *β ≥* 2, the dominant term in *g*(*ω*) is proportional to *ω*^*β*^ if and only if *b*_0_ = 1 and *b*_1_ = = *b*_*β/*2−1_ = 0 ≠*b*_*β/*2_.

*Proof*. We already established the base case *β* = 2. For the induction hypothesis, suppose that the statement holds for some *β* = *k*. We want to show that the statement holds for *β* = *k* + 2. If all terms in *g*(*ω*) up to order *k* − 2 vanish, then

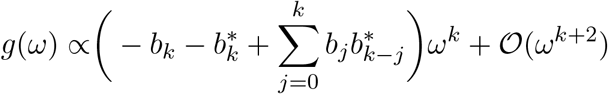

By the induction hypothesis, *b*_0_ = 1 and *b*_1_ = = *b*_*k/*2−1_ = 0, so this reduces to

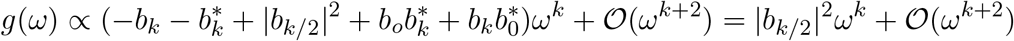

Hence the terms up to order *k* vanish if and only if *b*_0_ = 1 and *b*_1_ = *…* = *b*_(*k*+2)*/*2−1_ = 0. If, in addition, *b*_(*k*+4)*/*2−1_ = 0, then the order *k* + 2 term vanishes. We conclude that the dominant term in *g*(*ω*) is *ω*^*k*+2^ if and only if *b*_0_ = 1 and *b*_1_ = *…* = *b*_(*k*+2)*/*2−1_ = 0 *≠ b*_(*k*+4)*/*2−1_. That is, the statement holds for *β* = *k* + 2; by induction, it holds for all even *β ≥* 2. □

Next observe that

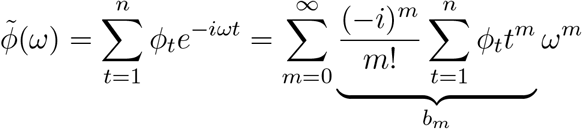

so we can rephrase Proposition S2.2.1 as follows: the dominant term in *g*(*ω*) is of order *β* if and only if *a*_0_ = 1 and *a*_1_ = *…* = *a*_*β/*2−1_ = 0 ≠*a*_*β/*2_, where 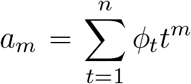. Let 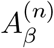 be the set of all order-*n* history kernels that satisfy these constraints (i.e. the basin of attraction of the *β* fixed point) and let

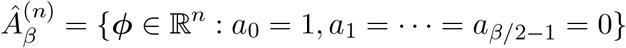

Then 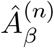 is an affine subspace of ℝ^*n*^ with dimension max(0, *n*−*β/*2) and 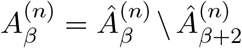. The nested structure of the constraints translates to 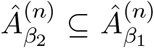 for all *β*_2_ *≥ β*_1_. For example, in the order-2 case, 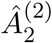 is the line through (0, 1) and 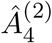 is the singleton {(2, −1)}, and 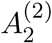 (the set order-2 AR models that flow into the *β* = 2 fixed point) is the line 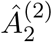 minus the point 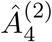.

A technical point is that, because the probability distribution of trajectories (Eq. (2)) that we used when performing tRG is only valid for stationary AR models, the actual tRG basins of attraction are the subsets of 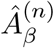 that are on the border between stationary and nonstationary AR models; this border consists of all the history kernels ***ϕ*** for which the smallest root of the characteristic polynomial 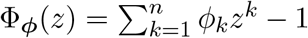 lies on the unit circle [1]. For example, 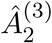 is a plane, but the set of AR(3) models that flows into the *β* = 2 fixed point is a triangular subset of this plane (Fig. 2F).

### 2.3. KL Divergence Rate for AR models

Fix two order-*n* AR models A and B with parameters (*ϕ*_A_, *σ*_A_) and (*ϕ*_B_, *σ*_B_), respectively. We seek an analytical expression for the KL divergence rate *J*(A *∥* B). Let 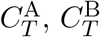 be the covariance matrices for a sequence of *T* steps **x** = (*x*_1_, …, *x*_*T*_)^*T*^ generated by each model. Then

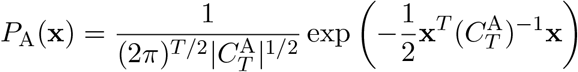

so that

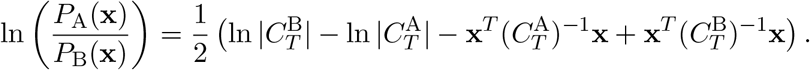

Since 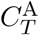 is symmetric and positive definite, its inverse is as well, and thus has a Cholesky decomposition:

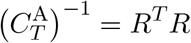

where *R*^*T*^ is lower-triangular. Making the change of variables **y** = *R***x** (equivalently **x** = *R*^−1^**y**), we get

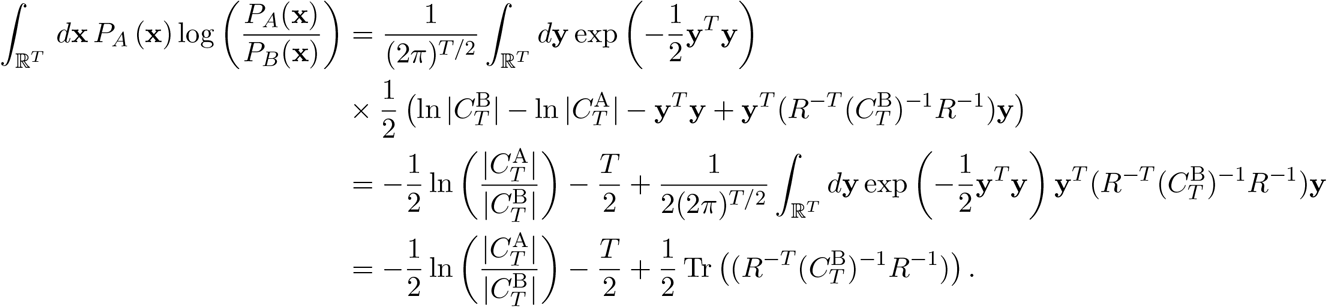

In the last step, we used the fact that for a real symmetric matrix *A*, if **y** *~ N* (0, *I*_*n*_), then *E*[**y**^*T*^ *A***y**] = Tr(*A*). See [2] for a similar derivation.

Since the trace of a matrix product is invariant to cyclic permutations,

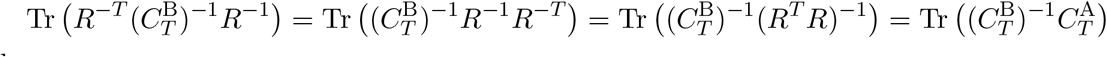

we have

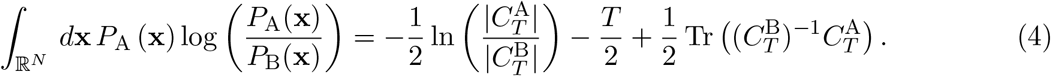

Next, note that the covariance matrices 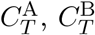 are Toeplitz; that is, for any *t, t*′ *∈* {1,, *T*}, we have 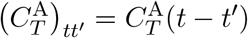 and similarly for B. Szego’s theorem [3] then tells us that

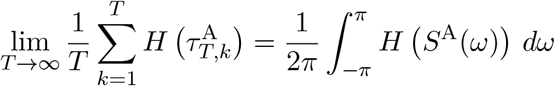

for any function *H* that is continuous on the range of *S*^A^(*ω*), where *S*^A^(*ω*) is the power spectral density of model A and 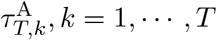 are the eigenvalues of 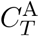.

Setting *H* to the identity function, we get

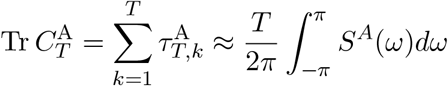

Similarly, setting *H*(*x*) = ln(*x*) yields

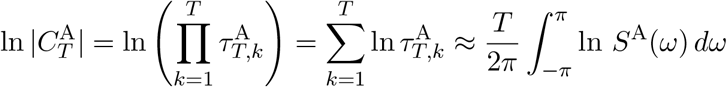

so that

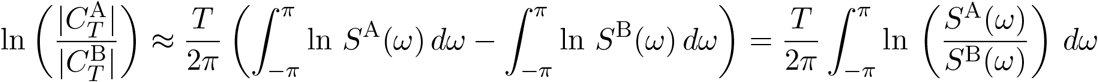

To simplify the trace term, 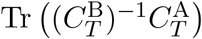, we use the fact that the sequence of the product of Toeplitz matrices 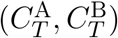 is asymptotically Toeplitz (see [3]), and that the spectral density of the product 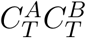 is (asymptotically) the product of the spectral densities *S*^*A*^ and *S*^*B*^; that is,

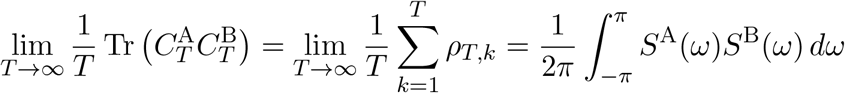

where *ρ*_*T,k*_ are the eigenvalues of the product 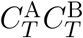. For the inverse of such matrices,

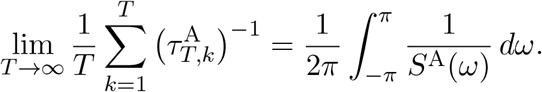

Putting these two facts together gives us

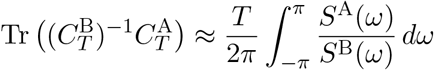

and thus

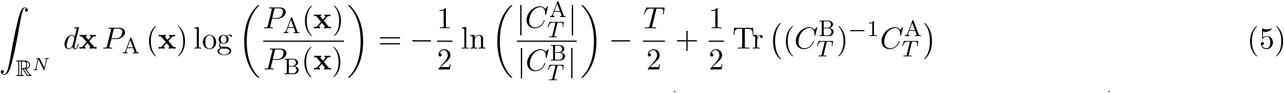

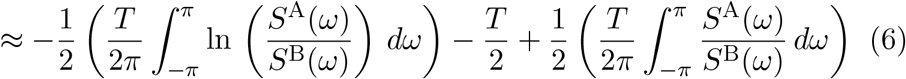

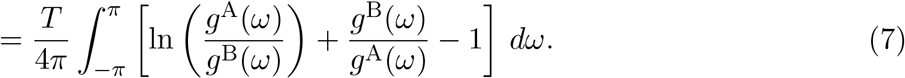

where 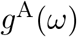 and 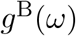 are defined as in Eq. (3). Rearranging the factor of *T* and taking the *T* → *∞* limit yields the formula for KL divergence rate in the main text.

### 2.4. Numerical Minimization of KL Divergence Rate

We have identified the tRG basins of attraction for AR models (Section S2.2) and derived a formula for the KL divergence rate between AR models (Section S2.3). To calculate *d*_*β*_ for a given AR model (call it A), it remains to minimize the KL rate *J*(A || B) over the set of all models B in the basin of attraction of the *β* fixed point. To do this, we used Matlab’s nonlinear constrained optimization solver fmincon. Specifically, we parameterized 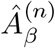 by the first *n*−*β/*2 coefficients of the history kernel (these are enough to uniquely identify a point on 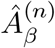 since it has dimensionality *n*−*β/*2). We initialized the optimization process at the point on 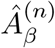 that was closest in Euclidean distance to the history kernel of model A. To ensure that the optimizer stayed on the critical manifold (which is a subset of 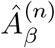; see S2.2), we used the inequality constraint

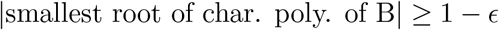

where *ϵ* = 10^−2^ is a numerical tolerance. For complete implementation details, see our publicly available toolkit [*github*…*doi*…*TBD*].

### 2.5. Further details on AR model estimation and *d*_2_ error bars

In the time-resolved *d*_2_ analysis in Fig. 4H, we fitted AR models to time series by maximum likelihood estimation. Here we detail this procedure in the general case of a segmented time series 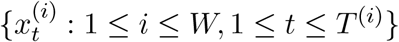 (here 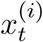 is the value of the time series at time step *t* during segment *i, W* is the number of segments, and *T* ^(*i*)^ is the duration of the *i*th segment), although in Fig. 4H the time series were not segmented (i.e. *W* = 1). For a given AR model order *n*, we maximized the log-likelihood

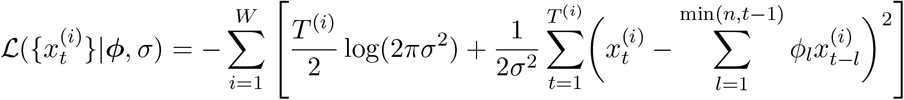

First, fix ***ϕ*** and find the value of *σ* (call it *σ*\*) that maximizes the likelihood:

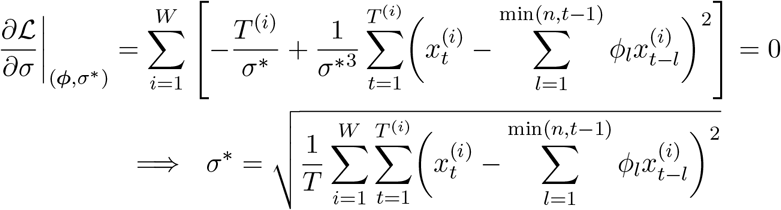

Here we introduced 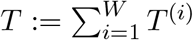. Substituting this back into the likelihood function yields

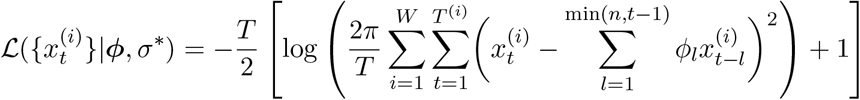

Maximizing this likelihood function is equivalent to minimizing the sum of squared errors. We do this in the usual way; specifically, for each *i* = 1, …, *W*, let *X*^(*i*)^ be the *T* ^(*i*)^ *×n* matrix with entries

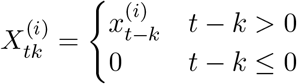

and let 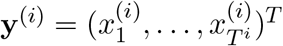. Then the maximum likelihood history kernel is

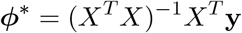

where

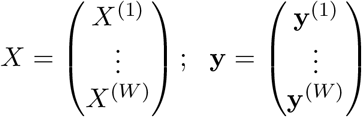

Next, to obtain uncertainties for the estimated history kernel, we compute the Hessian of the log-likelihood,

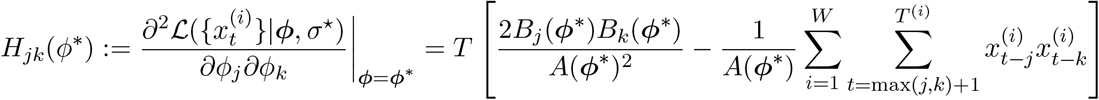

where

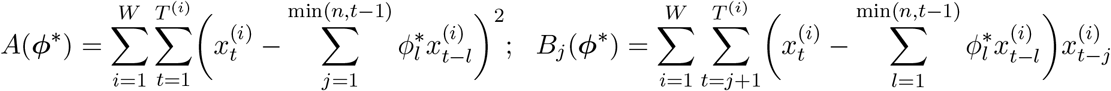

The matrix *H*(***ϕ***\*) encodes the quadratic approximation of the log-likelihood function around its minimum at ***ϕ***\*. Moreover, if we treat the likelihood (of the data, given a particular history kernel) as the likelihood of the kernel given the data (this is equivalent to adopting a uniform prior on the kernel), then the standard deviation of *d*_2_(***ϕ***) is

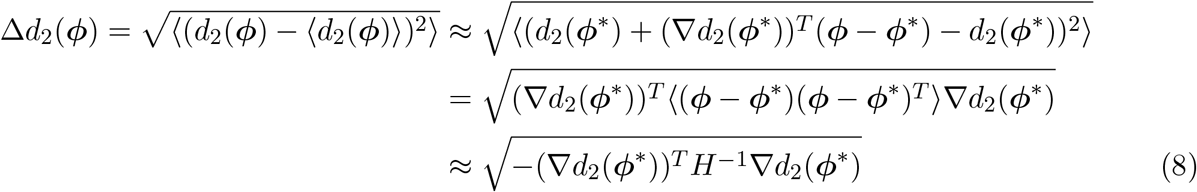

where ⟨·⟩ denotes an average over the likelihood of the kernel given the data. We used Eq. (8) to compute the *d*_2_ error bars shown in Fig. 4.

## 3. Model Benchmarking

In the main text Section II and Fig. 3, we showed that *d*_2_ faithfully represented the ground truth distance to criticality for three test case models. Here we detail this process more fully.

### 3.1. Numerical calculation of KL divergence rate

Much of this section relies on numerically calculating KL divergence rates between pairs of models, so we will first describe this procedure in general terms. Consider a family of time series models, with each model corresponding to a list of parameters *θ* and a conditional distribution *P*_*θ*_(*x*_*t*_|*x*_*t*−1_, *x*_*t*−2_, …) ≡ *P*_*θ*_(*x*_*t*_|past). We want to calculate the KL divergence rate *J*(*θ*_A_||*θ*_B_) between two models in this family with parameter values *θ*_A_ and *θ*_B_. In lieu of an analytical expression, we use the following numerical procedure:

(1) Run model A for *T* time steps by repeatedly sampling the distribution 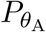 (*x*_*t*_|past).
(2) Repeat this *N* times to generate an ensemble of time series 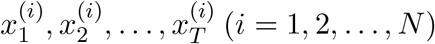, each of length *T*.
(3) Calculate the log-likelihoods

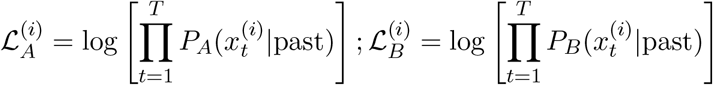

for each time series under models A and B.
(4) Estimate

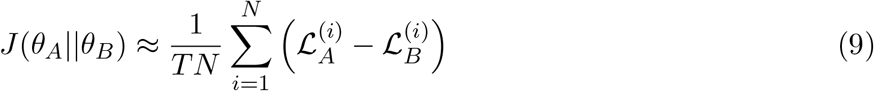

In the limit of large *T* and large *N*, this estimate becomes exact.

### 3.2. Univariate Hawkes Process

Since real neural activity is discretized into point events (“spikes”), we considered a univariate Hawkes process. In this model, spikes are modeled as a Poisson process with time-varying rate

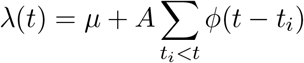

where the sum is over all spike times earlier than *t, µ* is a background spiking rate, and *ϕ*(*t*) is a history kernel (Fig. 3A). For simplicity, we choose *ϕ*(*t*) = *e*^−*t*^. Regardless of the value of *µ*, the model is stable for *A <* 1 and unstable for *A >* 1; at the transition point *A* = *A*_*c*_ = 1, the spiking statistics are scale-invariant [4], thus the model is at criticality.

Our definition of distance to criticality for time series models was applied here with some changes. First, imagine running a Hawkes process with parameters *A, µ* until it generates *K* spikes. Let the log-likelihood that these spikes occur at times *t*_1_, …, *t*_*K*_ be ℒ_*A,µ*_(*t*_1_, …, *t*_*K*_). (We derive a simple expression for this likelihood function in Prop. S3.2.1.) Then the KL divergence per unit time from the model *A, µ* to a critical model *A*_*c*_, *µ*_*c*_ is

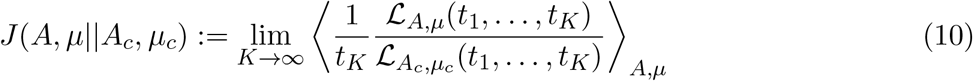

where ⟨·⟩ _*A,µ*_ denotes an average over realizations of the model with parameters *A, µ*. As before (cf main text Eq. (1)), distance to criticality is the minimum of this KL divergence rate taken over the set of critical models. In this case, this means minimizing over *µ*_*c*_. We calculated the expression in Eq. (10) with the numerical procedure described in Section S3.1. Specifically, we draw many sequences *t*_1_, …, *t*_*K*_ of spike times from the model with parameters *A, µ* using the algorithm of [5], calculate the quantity inside the angled brackets in Eq. (10) for each draw, and average over draws. We then perform a grid search for the optimal *µ*_*c*_ (Fig. 3C). The results of this calculation are the black points shown as a function of *A* in Fig. 3B.

Can an AR model-based approach recover the ground-truth distance to criticality? For a given realization of a Hawkes process, we counted the number of spikes in time bins of length Δ*t* and then fitted an AR model to the resulting spike-count time series. The best-fit AR model’s distance to criticality closely agrees with the ground-truth value obtained from Eq. (10) across a wide range of values of *A* (Fig. 3B,C).

#### Prop. S3.2.1

For the Hawkes process with parameters (*A, µ*), the probability distribution for the next spike time given the spiking history *t*_*i*−1_, *t*_*i*−2_, … is

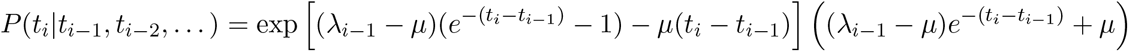

where *λ*_*i*−1_ is the rate immediately after the spike at time *t*_*i*−1_. The log-likelihood of a sequence of spike times (given an initial spike at *t*_0_ = 0) is then

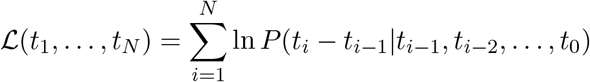

*Proof*. First, calculate the likelihood that the next spike occurs later than time *τ* after the most recent spike at time *t*_*i*−1_:

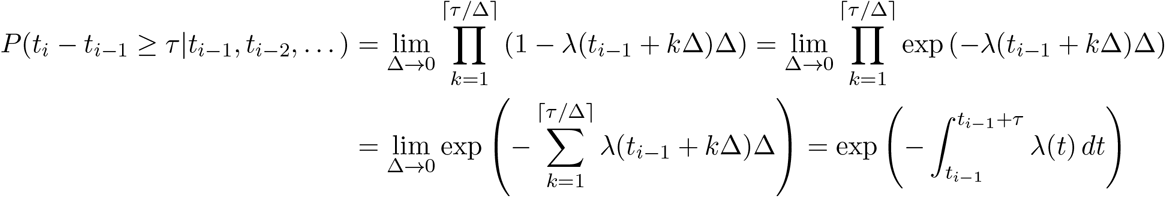

Here *λ*(*t*) is the rate at time *t* given that there were no spikes between *t*_*i*−1_ and *t*. It is easy to show (taking advantage of the property 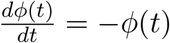 of the exponential kernel) that

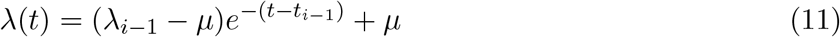

so

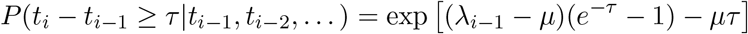

Differentiating with respect to *τ* then yields the desired expression. □

### 3.3. Nonlinear model: Langevin dynamics in quartic potential

Next, we tested our distance to criticality pipeline on a nonlinear time series model, the overdamped Langevin equation in a quartic potential:

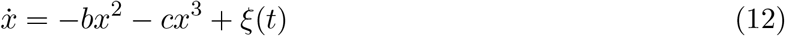

where as usual *ξ* is Gaussian white noise with ⟨*ξ*(*t*) ⟩ = 0 and ⟨*ξ*(*t*)*ξ*(*t*′)⟩ = *σ*^2^*δ*(*t* −*t*′). For what values of the parameters (*b, c, σ*) is this model at criticality? Since there is a transition from stable to unstable dynamics at *c* = 0 regardless of the values of *b* and *σ*, an initial guess might be that the half-plane of models {(*b, c, σ*): *c* = 0, *σ >* 0} are all at criticality. However, if *b* ≠ 0, then scale-invariance does *not* emerge as *c* → 0^+^. Instead, the model exhibits weak, short timescale fluctuations around the fixed point *x*\* = *b/c* of the deterministic dynamics. This can be seen by changing variables to *z* = *x* − *x*\*:

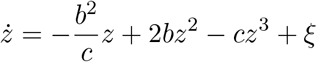

If *b* ≠ 0, then as *c* → 0^+^, the coefficient of the linear term approaches −*∞*, making the timescale of fluctuations in *z* approach zero. Hence the only critical models in this family are the continuous-time random walks ℳ_crit_ = {(*b, c, σ*): *b* = *c* = 0}.

To calculate the ground-truth distance to criticality for models in this family, we followed the same numerical strategy as for the Hawkes process, except with a time series instead of spikes. Specifically, we simulate Eq. (12) in discrete time steps of size Δ*t* and calculate the average (over many simulations) of the log-likelihood ratio between the true model (*b, c, σ*) and critical models (*b*_*c*_ = 0, *c*_*c*_ = 0, *σ*_*c*_), per unit time; we then choose the value of *σ*_*c*_ that minimizes this quantity. Repeating this procedure for various values of Δ*t* shows that the measured distance to criticality limits to a finite value as Δ*t* → 0. Moreover, applying the AR model pipeline to sample time series produced by the model Eq. (12) closely reproduces the ground-truth distance to criticality (Fig. 3E). This example illustrates the importance of a non-parametric definition of distance to criticality in systems with more than one relevant parameter, as it is not at all clear a priori how to combine *b* and *c* to measure deviation from the critical point at *b* = *c* = 0.

To calculate the KL divergence rate between a given model (*b, c, σ*) and the models in ℳ_crit_, we approximate Eq. (12) with the discrete update equation

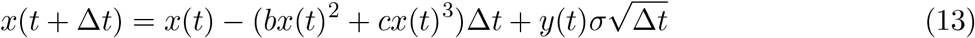

where *y*(*t*) ~ 𝒩 (0, 1) and is drawn independently for each *t* [cite]. The probability distribution for *x*(*t* + Δ*t*) given *x*(*t*) under the discrete dynamics Eq. (13) is

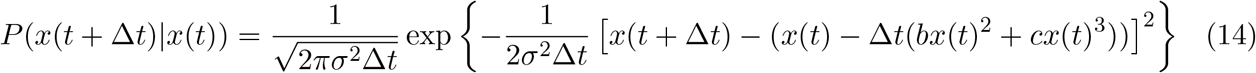

so the log-likelihood of a *T*-step discrete trajectory *x*(*t*_0_ + Δ*t*), *x*(*t*_0_ + 2Δ*t*), …, *x*(*t*_0_ + *T* Δ*t*) given the starting point *x*(*t*_0_) is

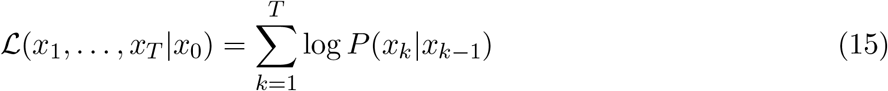

where for compactness we abbreviate *x*_*k*_:= *x*(*t*_0_ + *k*Δ*t*). To calculate the KL divergence rate from the model *A* = (*b, c, σ*) to a critical model *B* = (0, 0, *σ*_*c*_) *∈* ℳ _crit_, we follow the procedure described in Section 3. First, we draw *N* trajectories 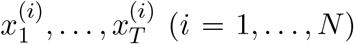 of duration *T* steps from model *A* with Eq. (13). Second, we use Eq. (15) to calculate the log-likelihoods 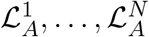 and 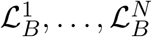 of these trajectories under *A* and *B*, respectively. Then Eq. (9) becomes

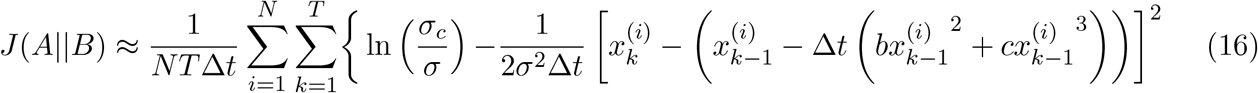

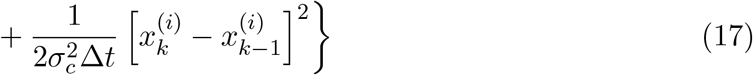

where we added the extra factor of 1*/*Δ*t* to put *J* in units of nats per time (rather than nats per time step). To calculate distance to criticality, we minimize *J*(*A*||*B*) over *σ*_*c*_:

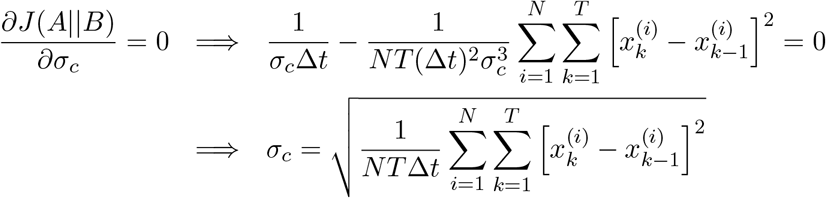

Plugging this expression into Eq. (17) then yields the distance to criticality.

### 3.4. Bursty model: Langevin dynamics with multiplicative noise

In addition to spiking and nonlinearity, another feature of neural systems that is absent in AR models is burstiness. In awake, adult animals, brain activity is not bursty, as we will show in the next section. However, in other situations, brain dynamics can be rather bursty. Examples include in vitro systems [6, 7, 8, 9], some anesthetized states [10], and newborn animals [11, 12]. AR models, on the other hand, produce time series that are symmetric about their mean and have no definite floor. As a third test case, we considered a simple model with multiplicative noise that captures both bursty and non-bursty dynamics,

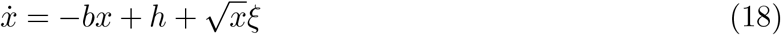

where as before *ξ* is Gaussian white noise with ⟨*ξ*(*t*) ⟩ = 0 and ⟨*ξ*(*t*)*ξ*(*t*′)⟩ = *σ*^2^*δ*(*t* −*t*′). When *h* (which can be interpreted as external drive) is close to zero, this model exhibits silence-separated bursts because of the 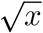 multiplicative noise. Increasing *h* moves the stable fixed point at *x* = *h/b* to larger *x*, making the dynamics less bursty. For any *h ≥* 0, the model transitions from stable to unstable dynamics and exhibits scale-invariance at *b* = 0. Hence the critical models in this family are those on the half-line *b* = 0, *h ≥* 0.

We calculated ground-truth distance to criticality in the same way as for the model Eq. (12), except that here there are two parameters (*h*_*c*_ and *σ*_*c*_) to vary when minimizing the KL divergence rate over the set of critical models (Fig. 3F). In brief, we simulated Eq. (18) in discrete time steps by sampling the exact solution to the associated Fokker-Planck equation, as described in [13]. (A naive Euler-type update equation similar to Eq. (13) is inadequate because of the 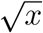 multiplicative noise.) We then again used the solution to the Fokker-Planck equation given in [13] to compute the likelihood of the simulated trajectories under the actual model (parameters (*b, h, σ*)) and a critical model (parameters (*b*_*c*_ = 0, *h*_*c*_, *σ*_*c*_)). Next, we plugged the result into (9) to compute the KL divergence rate from the model (*b, h, σ*) to the critical model (*b*_*c*_ = 0, *h*_*c*_, *σ*_*c*_). Finally, we performed a grid search to find the *h*_*c*_ and *σ*_*c*_ that minimized the KL rate.

Comparing the ground-truth distance to criticality to that obtained via AR model fits reveals close agreement across a large swath of values of *b* and *h* (Fig. 3G). This agreement breaks down as *h* → 0^+^ and the dynamics become more bursty. Put simply, our approach is robust to low levels of burstiness, but breaks down for very bursty data.

**Figure 5.**
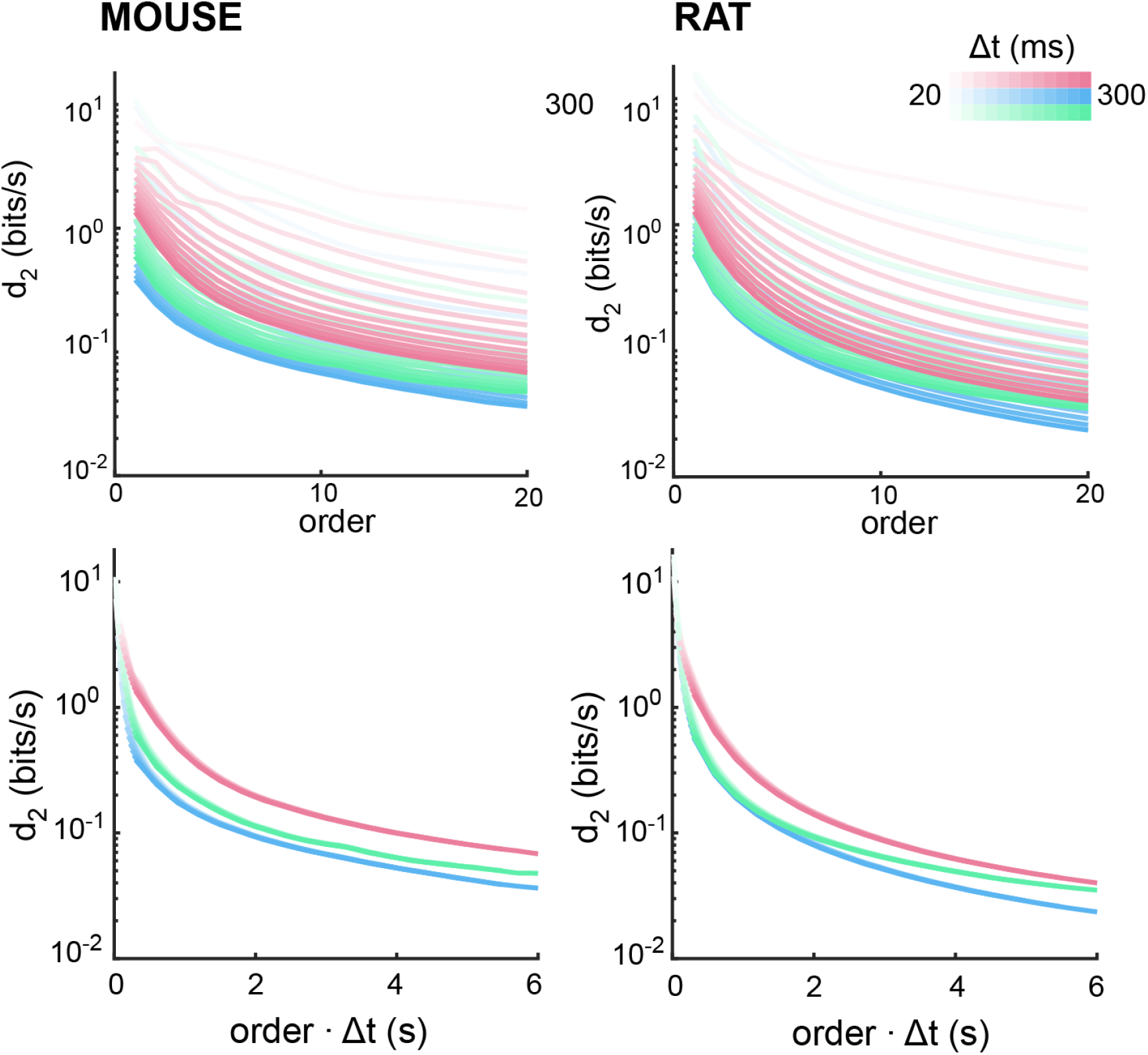
Temporal reach explains dependence of *d*_2_ on time bin and model order. In general, when applying our approach to spike data, *d*_2_ will depend strongly on the choice of time bin Δ*t* used to count spikes and the choice of AR model order. The top row shows how *d*_2_ varies with model order and Δ*t* for the mouse data (left) and rat data (right) and each sleep/wake state (blue - wake, red - NREM, green - REM). The wide ranging *d*_2_ values collapse onto a single function when considered in terms of “temporal reach”, i.e. the product of (model order) *×* Δ*t*. The temporal reach is the time (in seconds) that the AR model kernel reaches into the past.

**Figure 6.**
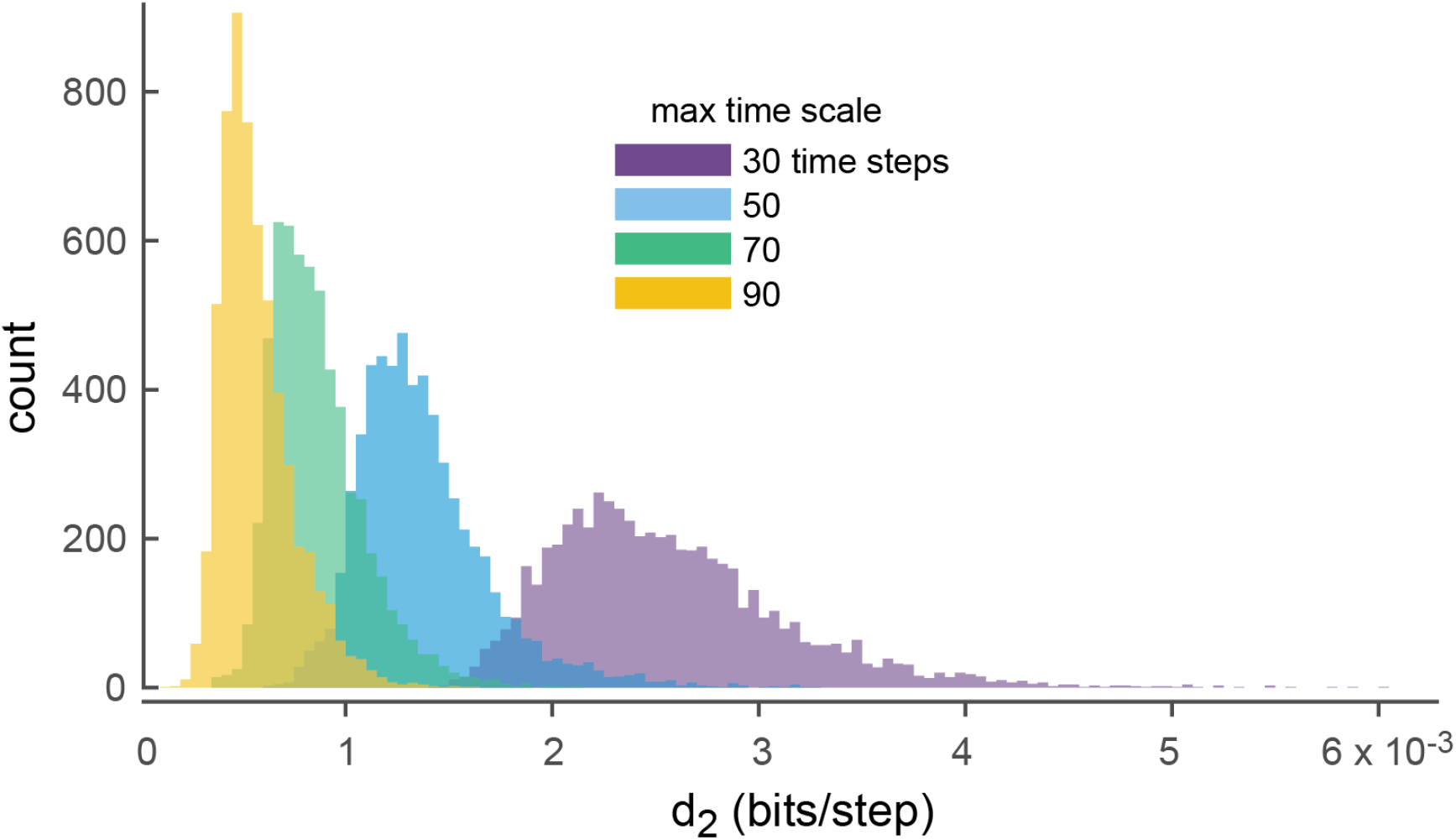
Distance to criticality (*d*_2_) is not solely determined by the longest time scales of a system. We considered four ensembles of AR(5) models, each with a fixed maximum time scale (color indicates max time scale). For fixed max time scale, distance to criticality *d*_2_ can vary by as much as an order of magnitude. Here, *d*_2_ was calculated analytically from the AR history kernel. Thus, the variability in *d*_2_ for fixed time scale is not due to statistical noise in simulations.

## 4. Dependence on Time Bin Length and AR Model Order

When working with spike data, how should one choose the time bin Δ*t* for counting spikes and the model order for calculating *d*_2_? These are the two data- analytic parameter choices that most impact the values of *d*_2_. One important fact that helps clarify this choice is demonstrated in Supplementary Fig. 5. We show that two *d*_2_ measurements are comparable if model order and *×* Δ*t* are chosen such that their product is fixed. The product (model order) × Δ*t* is the “temporal reach” (in seconds) of the AR model history kernel. If this temporal reach is the same for two different *d*_2_ measurements, they can be fairly compared. Otherwise, comparisons should not be made.

## 5. Relating *d*_2_ to the longest time scale of the dynamics

Many previously established data analytic methods assess the dominant time scale of a time series. These include estimates of the autocorrelation time [14, 15, 16] and other methods. Such methods are expected to be related to proximity to criticality because dominant timescales diverge as a system approaches criticality - a phenomenon sometimes called ‘critical slowing down’. Thus, we expect *d*_2_ to be related to the dominant time scale. However, we show in Supplementary Fig. 6 that true proximity to criticality (i.e. *d*_2_) is not simply a readout of the longest time scale of a system. We randomly generated many AR(5) models all with the same maximum time scale. Here the maximum time scale is set by the root of the characteristic polynomial of the AR model that is closest to the unity circle. We found that the variability in *d*_2_ for fixed maximum time scale was comparable to the difference in average *d*_2_ caused by increasing the maximum time scale by 20 time steps. Thus, we conclude that the longest time scale does not determine proximity to criticality, at least in AR models and suggests that *d*_2_ will better capture proximity to criticality than methods that rely solely on measuring the dominant time scale.

## 6. Restricting the set of critical models to its intersection with a tractable family

Directly implementing our definition of distance to criticality (main text Eq. (1)) would require knowing the structure of the abstract set ℳ_*c*_ of *all* critical models. As a practical compromise, we restricted our attention (main text Section I.C) to the critical models lying within a tractable family of models, AR models. In doing so, we assumed that the nearest critical model (in the sense of KL rate; call this model B*) to the model whose DTC we are computing (call it A) can be approximated by a critical AR model; that is, B* *≈* B for some critical AR model B. More work is needed to understand when this approximation is valid; intuitively, however, if A is well-approximated by an AR model, then B* should be well-approximated by a critical AR model. We applied the same logic to the familes of models used for benchmarking (main text Section II; Section S3); when calculating the “ground truth” distance to criticality of a model in one of these families (e.g. Hawkes processes), we limited the KL rate minimization to critical models lying in the same family. This is admittedly pragmatic and not formally justified. A more flexible implementation of our definition of distance to criticality that does not rely on any specific class of models will be a target of future work.

